# Branched-chain amino acid fermentation as an alternative mammalian electron sink

**DOI:** 10.64898/2026.08.24.746737

**Authors:** Ayush D. Midha, Brandon T. L. Chew, Yolanda Martí-Mateos, Skyler Y. Blume, Will R. Flanigan, Brandon R. Desousa, Augustinus G. Haribowo, Aunoy Poddar, Saahil Chadha, Bruno B. Queliconi, Alec M. Barrios, Michela Traglia, Reuben Thomas, Junji Suzuki, Mito Kuroda, Steven J. Altschuler, Lani F. Wu, Yuriy Kirichok, Mercedes F. Paredes, Tracy G. Anthony, Polina V. Lishko, Isha H. Jain

## Abstract

Hypoxia disrupts mitochondrial respiration and increases the NADH/NAD^+^ ratio, causing reductive stress. To maintain redox homeostasis, mammalian cells divert electrons toward fermentation. While fermentation in mammals typically involves lactate production, we identify the fermentation of branched-chain amino acids (BCAAs) as an alternative electron sink activated by hypoxia. The resulting metabolites are excreted in urine as a distinct mechanism for alleviating reductive stress. BCAA fermentation is catalyzed by lactate dehydrogenase (LDH) enzymes and is highly responsive to the NADH/NAD^+^ ratio. Consequently, BCAA fermentation products are sensitive biomarkers for reductive stress in human contexts ranging from resistance exercise to severe hypoxemia. Furthermore, we find that mouse sperm have evolved highly efficient BCAA fermentation, providing a specific metabolic strategy to support the anaerobic electron flow that facilitates flagellar hypermotility across mammalian sperm. Our work highlights an under-appreciated fate of BCAAs in response to reductive stress.

## INTRODUCTION

Oxygen is the primary terminal electron acceptor for the mammalian electron transport chain (ETC). In the ETC, electrons from NADH are ultimately transferred to oxygen, and this process supports the bioenergetic^1^ and biosynthetic^2,3^ demands of the cell. Impaired respiration slows NADH utilization and increases the NADH/NAD^+^ ratio—a condition termed reductive stress,^4–6^ which contributes to the pathophysiology of diverse metabolic diseases, including diabetes,^6–8^ fatty liver disease,^9–11^ and mitochondrial disease.^12^

In low-oxygen conditions, cells use alternative electron acceptors to regenerate NAD^+^. For example, cells can divert electrons to fumarate within the ETC.^13,14^ Alternatively, cells can use fermentation, which involves NADH oxidation independent of the ETC.^15^ In animals, the most common form of fermentation is anaerobic glycolysis. During anaerobic glycolysis, pyruvate generated from glucose is reduced to lactate instead of being further oxidized to enter the tricarboxylic acid (TCA) cycle. In hypoxic cells, this is induced by the hypoxia-inducible transcription factors (HIF).^16,17^ In normoxic conditions, HIFs are hydroxylated by prolyl hydroxylase (PHD) enzymes in an oxygen-dependent reaction, marking them for degradation. In hypoxia, the un-marked HIF proteins translocate to the nucleus, complex with the Aryl Hydrocarbon Receptor Nuclear Translocator (ARNT), and induce the expression of genes required for adaptation to hypoxia, including enzymes involved in anaerobic glycolysis.^18–20^

While HIF-induced lactate production is ubiquitous across the animal kingdom, other fermentation pathways have also been characterized. For example, anoxia-tolerant goldfish ferment glucose to ethanol, which is then excreted.^21–23^ In anaerobic oyster hearts, the fermentation of aspartate regenerates NAD^+^ to support glycolysis.^24,25^ Similar hypoxia-induced pathways have been identified as electron sinks in mammalian cells, including proline synthesis,^26^ fatty acid desaturation,^8^ and the production of the α-hydroxyacids α-hydroxyglutarate^27–29^ and α-hydroxybutyrate.^2,11^ However, the tissue-specific functional relevance of these alternative electron acceptors remains unknown.

Here, we identify branched-chain amino acid (BCAA) fermentation as an alternative electron sink in mammals. Unlike lactate, which is preserved within circulation as a fuel source,^30,31^ the products of BCAA fermentation are primarily excreted, functioning as a disposal mechanism for excess reducing equivalents. We find that these metabolites, which have previously been studied in fermentative micro-organisms, are produced in mammalian cells by the catalytic promiscuity of lactate dehydrogenase enzymes and are highly responsive to reductive stress. Furthermore, we show that mouse sperm have evolved a uniquely efficient capacity to ferment BCAAs. This represents a specific metabolic solution to the problem of anaerobic electron flow, which is required for flagellar hypermotility in all mammalian sperm. Collectively, these findings highlight the diversity of anaerobic metabolism in mammals and define BCAA fermentation as a conserved response to reductive stress.

## RESULTS

### Hypoxia and HIF activation promote BCAA fermentation

To systematically characterize metabolic adaptations to hypoxia, we profiled the metabolites in organs harvested from mice continuously exposed to 21% O_2_ (normoxia), 11% O_2_ (moderate hypoxia), and 8% O_2_ (hypoxia) or treated with the PHD inhibitor FG-4592, which results in increased HIF stabilization in normoxia (**Figure 1A**). We collected plasma and 6 organs (heart, white adipose tissue, brain, lung, muscle, and liver) from mice at acute (3 hours, 24 hours) and chronic (1 week, 3 weeks) time points in hypoxia. The resulting metabolomics data reveal the time- and organ-specific metabolic adaptations to hypoxia, and these data are publicly available at https://jain-lab-ucsf.github.io/hypoxia-metabolomics/. While organs exhibited unique metabolic signatures throughout the course of hypoxia (**Figure S1A-C**), we identified a core set of metabolites that were commonly enriched across multiple tissues (**Figure S1D**). Leveraging this dataset, we stratified metabolic alterations into HIF-dependent and HIF-independent responses (**Figure S1E**) and classified metabolites as responsive to acute versus chronic hypoxia (**Figure S1F**).

**Figure 1.**
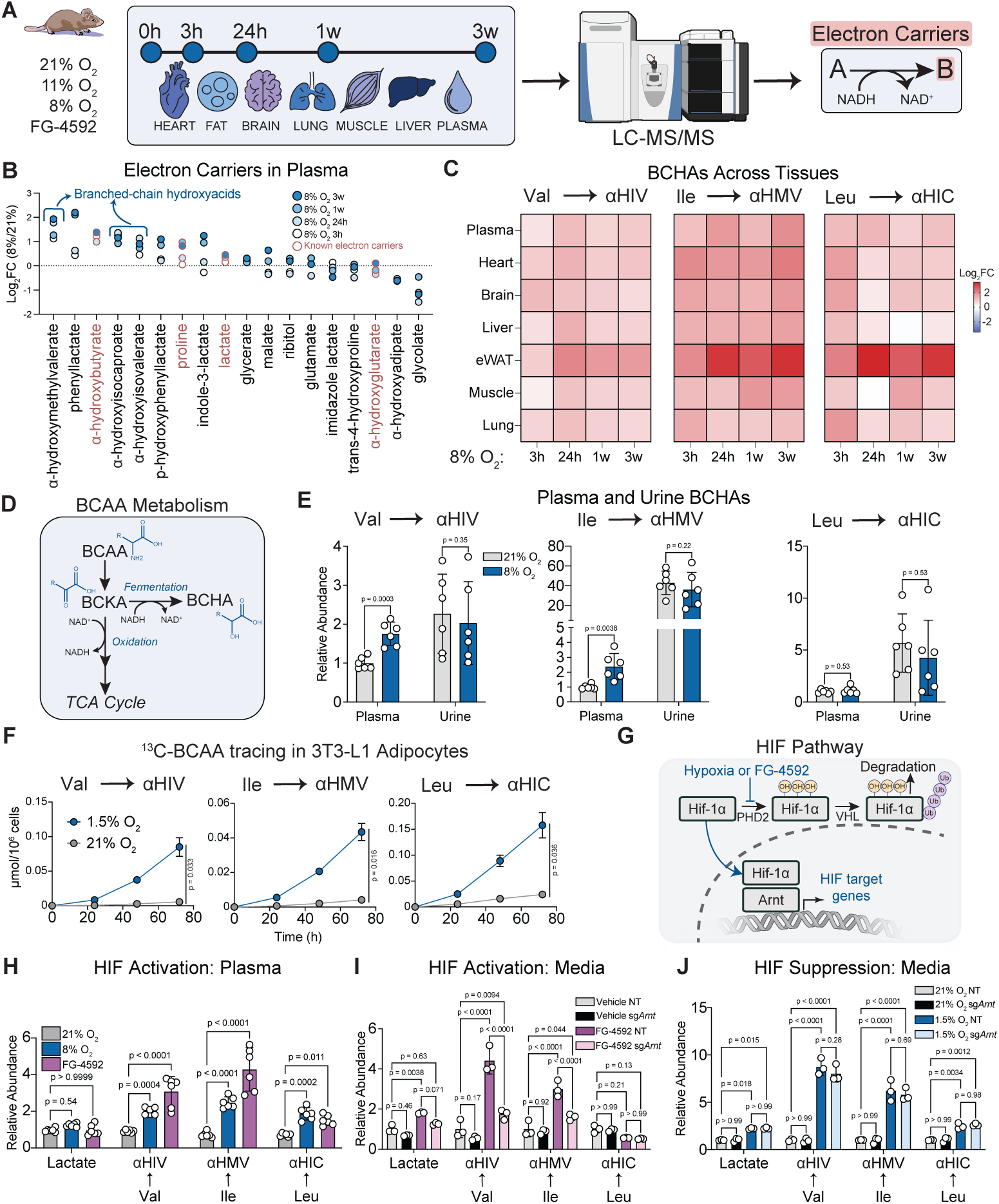
Hypoxia and HIF activation promote BCAA fermentation. (A) Schematic depicting the use of mouse tissue metabolomics in hypoxia to identify putative electron carriers. (B) Fold change enrichment of electron carriers in plasma from mice housed at 8% O_2_ for 3 hours, 24 hours, 1 week, or 3 weeks compared to mice at 21% O_2_. n = 6 mice per group. (C) Fold change enrichment of BCHAs in all sampled tissues from mice housed at 8% O_2_ for 3 hours, 24 hours, 1 week, or 3 weeks compared to mice at 21% O_2_. (D) Schematic depicting the oxidative and fermentative catabolism pathways for BCAAs. (E) Concentration of α-hydroxyisovaleric acid in plasma and urine from mice housed at 21% O_2_ or 8% O_2_ for 24 hours. Plasma 21% n = 4 mice, plasma 8% n = 6 mice, urine 21% and 8% n = 5 mice. (F) Concentration of ^13^C-BCHAs in conditioned media from 3T3-L1 adipocytes cultured at 21% O_2_ or 1.5% O_2_ for 24, 48, and 72 hours. n = 3 wells. (G) Schematic showing the HIF pathway. (H) Volume-normalized relative abundance of lactate and BCHAs in plasma from mice housed at 21% O_2_ or 8% O_2_ and from mice treated with 100 mg/kg of FG-4592 twice 12 hours apart. n = 6 mice per group. (I-J) Relative abundance of ^13^C-BCHAs in conditioned media from control or *Arnt*-deficient 3T3-L1 adipocytes after 3-day treatment with 0.15% DMSO or 75 µM FG-4592 (I) or 3-day treatment with 21% O_2_ or 1.5% O_2_ (J). Mean ± SD are shown. Significance was determined by multiple two-sample t-tests with FDR (E), two-way ANOVA followed by Šídák’s multiple comparisons test (F) or Tukey’s multiple comparisons test (H, I, J).

Using these metabolomics data, we interrogated mammalian fermentation pathways by identifying putative anaerobic electron carriers. For this purpose, we defined anaerobic electron carriers as the products of reactions that consume NADH and produce NAD^+^. These reactions were compiled using the KEGG database of metabolic pathways^32^ with further manual additions and curation based on the literature. Many well-known electron carriers were enriched in plasma at certain timepoints in 8% O_2_, including lactate, proline,^26^ and α-hydroxybutyrate (**Figure 1B**).^11,33^ We also identified three metabolites that had not been extensively characterized in mammals— α-hydroxymethylvalerate (αHMV), α-hydroxyisocaproate (αHIC), and α-hydroxyisovalerate (αHIV). These metabolites, collectively named branched-chain hydroxyacids (BCHAs) were enriched in plasma at every time point during 8% O_2_ exposure (**Figure 1B**). BCHAs were also enriched in every profiled organ, with the greatest enrichment observed in epididymal white adipose tissue (eWAT) (**Figure 1C**, **S2A**).

BCHAs are derived from the BCAAs, valine, leucine, and isoleucine. Canonically, mammalian catabolism of BCAAs begins with their deamination by the enzyme branched-chain aminotransferase (BCAT) to form branched-chain keto-acids (BCKAs), which are subsequently oxidized by mitochondrial branched-chain keto-acid dehydrogenase (BCKDH) in an NADH-generating reaction. Just as pyruvate can be diverted from mitochondrial oxidation to form lactate in hypoxia, BCKAs can be shunted into an alternative fermentative pathway. This pathway, which is known in some bacteria^34–36^ but not understood in mammals, reduces BCKAs to form BCHAs and regenerate NAD^+^ (**Figure 1D**).^37^ Supporting the tradeoff between BCAA oxidation and fermentation, BCHAs have previously been identified in the urine of patients with maple syrup urine disease,^38^ a condition characterized by BCKDH deficiency. Based on this observation, we suspected that BCHAs may serve as a mechanism for excreting excess reducing equivalents. Indeed, we observed a high abundance of BCHAs in urine (**Figure 1E**), while lactate was undetectable in urine.

Because enzymes catalyzing BCHA production have been identified in fermentative bacteria (e.g., *Lactobacillus*,^34^ *Weissella*,^35^ and *Clostridioides*^36^ spp.), prior detection of BCHAs in mammals has been attributed to microbiome activity.^39^ To determine whether mammalian cells are capable of autonomous BCAA fermentation, we measured BCHA production from mammalian cells in culture. Given the substantial enrichment of BCHAs in eWAT and the established role of BCAA catabolism in adipocyte homeostasis,^40,41^ we differentiated 3T3-L1 murine fibroblasts into adipocytes for use as a model *in vitro* system. We cultured adipocytes in 21% O_2_ or 1.5% O_2_ in the presence of ^13^C-labeled BCAAs. After 72 hours, the conditioned media from hypoxic adipocytes exhibited higher concentrations of labeled BCHAs compared to normoxic controls (**Figure 1F**, **S3A**), confirming that murine adipocytes directly ferment BCAAs in hypoxia.

Next, we interrogated the regulatory role of HIF signaling in BCAA fermentation. Acute *in vivo* administration of the PHD inhibitor FG-4592, which stabilizes HIF (**Figure 1G**), elevated BCHA levels by up to 4-fold (**Figure 1H**). Similarly, in cultured adipocytes, 72-hour treatment with FG-4592 increased the fermentation of valine and isoleucine, an effect that was substantially attenuated in cells lacking the HIF interaction partner, ARNT (**Figure 1I**, **S3B**). In hypoxic cells, however, *Arnt* ablation did not suppress BCAA fermentation (**Figure 1J**, **S3C**). Taken together, these data indicate that HIF activation is sufficient, but not necessary, for BCAA fermentation, suggesting that other aspects of the cellular hypoxia response activate this pathway.

### LDHA catalyzes the fermentation of valine and isoleucine

We then set out to nominate candidate mammalian enzymes responsible for the fermentation of BCAAs. To this end, we leveraged a genome-wide association study of human serum metabolites in a cohort of Finnish men.^42^ Within this dataset, we observed that the circulating abundances of αHIV (derived from valine) and αHMV (derived from isoleucine) were significantly associated with single-nucleotide polymorphisms (SNPs) upstream of *LDHA*, a HIF target gene that encodes lactate dehydrogenase A. In contrast, levels of αHIC (derived from leucine) exhibited a much weaker association with this locus (**Figure 2A**). Both BCKAs and pyruvate, the canonical substrate of LDHA, are α-ketoacids. Moreover, prior work has demonstrated that the SNPs associated with αHIV levels cluster in a region that physically interacts with the *LDHA* promoter.^43^ Consistent with this link, we found that one of the SNPs associated with decreased levels of αHIV and αHMV is also associated with significantly lower *LDHA* mRNA expression in skeletal muscle (**Figure 2A**).^44^

**Figure 2.**
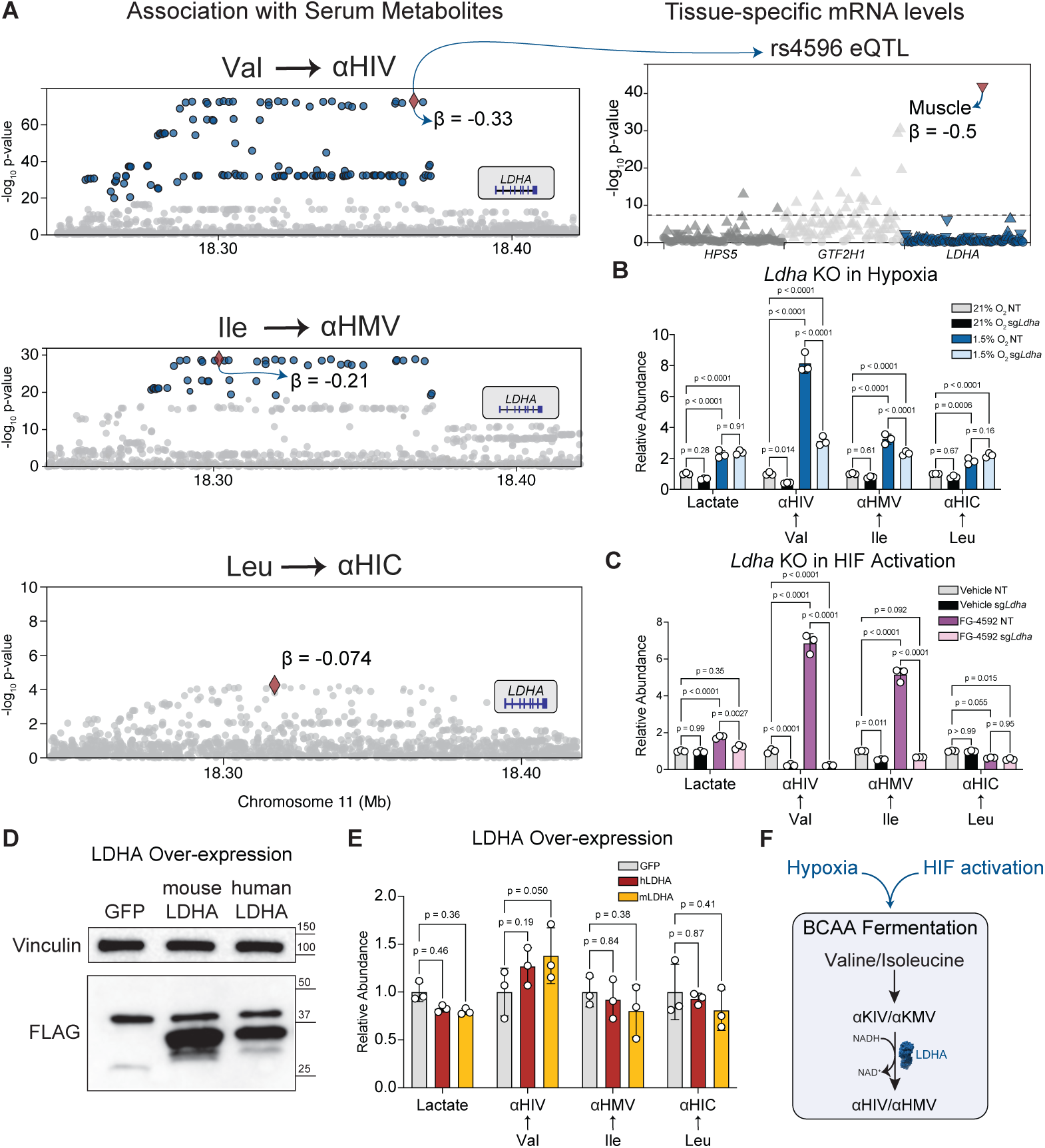
LDHA catalyzes the fermentation of valine and isoleucine. (A) Locus plot of single-nucleotide polymorphisms (SNP) upstream of *LDHA* showing the magnitude of association with branched-chain hydroxyacids from the METSIM cohort (Yin et al., 2022) (left) and expression quantitative trait locus (eQTL) plot showing the strength of association of one SNP with expression of *LDHA* in various tissues (right). Leading SNPs are depicted as diamonds, and all SNPs with p-values less than 10^-20^ are colored in blue. On the eQTL plot, triangles pointing up indicate a positive β and triangles pointing down indicate a negative β. (B-C) Relative abundance of lactate and ^13^C-BCHAs in conditioned media from control or *Ldha*-deficient 3T3-L1 adipocytes after 3-day treatment with 21% O_2_ or 1.5% O_2_ (B) or 3-day treatment with 0.15% DMSO or 75 µM FG-4592 (C). n = 3 wells. (D) Immunoblots of undifferentiated 3T3-L1 cells overexpressing GFP, mouse LDHA, or human LDHA. (E) Relative abundance of lactate and ^13^C-BCHAs in conditioned media from 3T3-L1 adipocytes overexpressing GFP, mouse LDHA (mLDHA), or human LDHA (hLDHA) after 3 days of incubation at 21% O_2_. n = 3 wells. (F) Schematic showing that hypoxia and HIF promote the fermentation of valine and isoleucine. Mean ± SD are shown. Significance was determined by two-way ANOVA followed by Tukey’s multiple comparisons test.

To directly test whether lactate dehydrogenase A is necessary for BCAA fermentation in hypoxia, we generated *Ldha*-deficient 3T3-L1 adipocytes. Deletion of *Ldha* significantly blunted the fermentation of valine and isoleucine, but not leucine, in cells exposed to hypoxia (**Figure 2B**, **S4A**), as well as in cells treated with FG-4592 (**Figure 2C**, **S4B**). *Ldha* ablation had a much smaller effect on lactate levels, likely because LDHB can also catalyze lactate production. Next, we asked whether increasing enzyme availability could drive BCAA fermentation. Heterologous expression of the *Weissella confusa* L-hydroxyisocaproate dehydrogenase, which is known to catalyze BCHA production, increased BCAA fermentation 60- to 250-fold (**Figure S4C**), but over-expression of mouse or human LDHA was not sufficient to promote BCAA fermentation (**Figure 2D, 2E**, **S4D**).

To explain this discrepancy, we measured the activity of human LDHA using αKIV (derived from valine) as a substrate. We observed that human LDHA exhibits a relatively high K_m_ for BCKAs (in the millimolar range) (**Figure S4E**). These kinetic parameters suggest that BCAA fermentation flux is governed largely by substrate abundance rather than enzyme abundance. Along these lines, we observed that supplementing wild-type cells with α-ketoisovalerate (αKIV, derived from valine) increased the production of its fermentation product αHIV by approximately 40-fold (**Figure S4F**). Collectively, these findings indicate that hypoxia and HIF activation promote the fermentation of valine and isoleucine via a mechanism that is catalytically dependent on, though not rate-limited by, lactate dehydrogenase A.

### Cytosolic NADH reductive stress drives valine and isoleucine fermentation

Because BCAA fermentation consumes NADH and regenerates NAD^+^, we hypothesized that an increase in the NADH/NAD^+^ ratio would activate BCAA fermentation, even in the absence of hypoxia. To test this hypothesis, we treated 3T3-L1 adipocytes with rotenone, an inhibitor of complex I of the electron transport chain (ETC). Complex I directly oxidizes NADH, so rotenone treatment increases the NADH/NAD^+^ ratio without inducing hypoxia. We found that rotenone was sufficient to increase lactate production and stimulate the fermentation of valine and isoleucine (**Figure 3A**, **S5A**).

**Figure 3.**
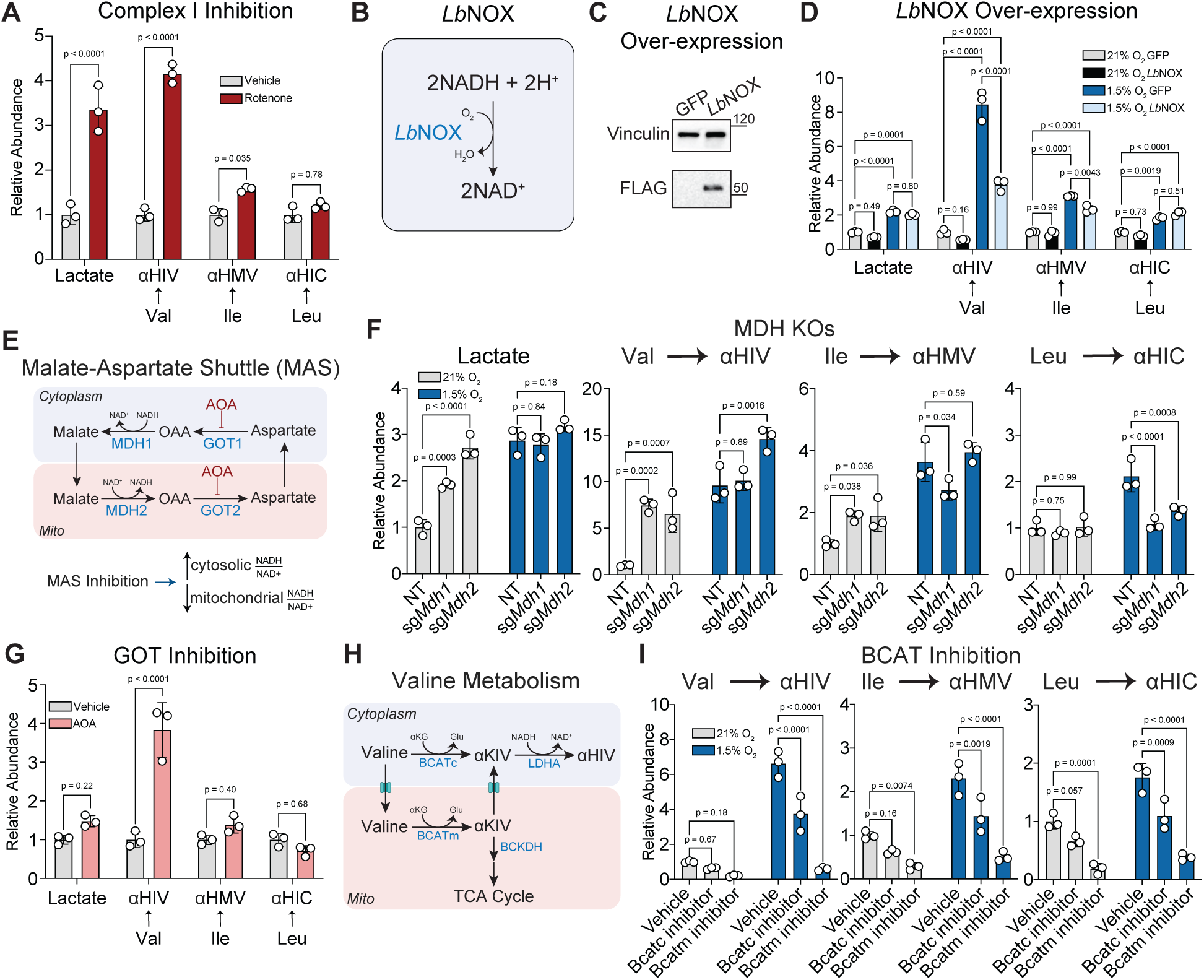
Cytosolic NADH reductive stress drives valine and isoleucine fermentation. (A) Relative abundance of lactate and ^13^C-BCHAs in conditioned media from 3T3-L1 adipocytes treated with 0.1% ethanol or 500 nM rotenone for 3 days. n = 3 wells. (B) Schematic depicting the reaction catalyzed by LbNOX. (C) Immunoblots of 3T3-L1 cells overexpressing GFP or LbNOX. (D) Relative abundance of lactate and ^13^C-BCHAs in conditioned media from 3T3-L1 adipocytes after 3-day treatment with 21% O_2_ or 1.5% O_2_. Cells were expressing *Lb*NOX. n = 3 wells. (E) Schematic depicting the malate-aspartate shuttle. (F) Relative abundance of lactate and ^13^C-BCHAs in conditioned media from control or *Mdh1*-deficient or *Mdh2*-deficient 3T3-L1 adipocytes after 3-day treatment with 21% O_2_ or 1.5% O_2_. n = 3 wells. (G) Relative abundance of lactate and ^13^C-BCHAs in conditioned media from 3T3-L1 adipocytes treated with 400 µM AOA for 3 days. n = 3 wells. (H) Schematic depicting the compartmentalized pathways in valine catabolism. (I) Relative abundance of lactate and ^13^C-BCHAs in conditioned media from 3T3-L1 adipocytes after 3-day incubation in 21% O_2_ or 1.5% O_2_ and treatment with 10 µM BCATc inhibitor 2 or 50 µM BCAT-IN-2 (BCATm inhibitor). n = 3 wells. Mean ± SD are shown. Significance was determined by two-way ANOVA followed by Šídák’s multiple comparisons test (A, G) or Tukey’s multiple comparisons test (D, F, I).

Conversely, to determine if an increase in the NADH/NAD^+^ ratio is necessary for BCAA fermentation, we overexpressed the *Lactobacillus brevis* NADH oxidase *Lb*NOX, which directly oxidizes NADH independently of the ETC (**Figure 3B, 3C**).^45^ Expression of *Lb*NOX in the cytosol significantly decreased valine and isoleucine fermentation in hypoxia (**Figure 3D**), confirming that NADH reductive stress is required for BCHA production in hypoxia.

We next sought to define the subcellular compartment where redox state controls fermentation. To do this, we manipulated the malate-aspartate shuttle (MAS), the primary mechanism for transferring reducing equivalents from the cytosol to mitochondrial matrix.^46^ Inhibition of the MAS elevates the cytosolic NADH/NAD^+^ ratio while simultaneously decreasing the mitochondrial NADH/NAD^+^ ratio (**Figure 3E**).^47^

We genetically disrupted the MAS by generating 3T3-L1 cells deficient in either *Mdh1* or *Mdh2* (**Figure 3E**). Strikingly, ablation of either enzyme was sufficient to increase valine and isoleucine fermentation in normoxia (**Figure 3F**, **S5C**). Interestingly, knocking out *Mdh1* or *Mdh2* completely reversed the hypoxia-induced increase in leucine fermentation in hypoxia, suggesting a distinct regulatory mechanism for the catabolism of leucine in hypoxia (**Figure 3F**, **S5C**). We corroborated our findings using an orthogonal pharmacological approach: treatment with aminooxyacetate (AOA), a small-molecule inhibitor of the MAS enzymes GOT1 and GOT2 (**Figure 3E**), similarly increased valine fermentation (**Figure 3G**, **S5D**). These data indicate that valine and isoleucine fermentation is responsive specifically to the cytosolic NADH/NAD^+^ ratio.

The catabolism of BCAAs begins with their deamination to BCKAs by BCAT, which exists as distinct mitochondrial (BCATm) and cytosolic (BCATc) isoforms.^37^ BCATm is expressed ubiquitously; BCATc expression is thought to be restricted to the brain,^37^ but there is evidence of cytosolic BCAT activity in other cell types, including brown adipocytes.^48^ To determine which pool of BCAT generates fermentable BCKAs (**Figure 3H**), we treated cells with isoform-specific inhibitors. Inhibition of BCATm almost completely abolished BCAA fermentation, whereas inhibition of BCATc had a significantly weaker effect (**Figure 3I**, **S5E**). These results indicate that during valine and isoleucine fermentation, BCKAs produced in mitochondria are exported to the cytosol for their reduction to BCHAs.

### Restrained BCAA oxidation is required for BCAA fermentation *in vivo*

Because a single pool of mitochondrially generated BCKAs fuels both the oxidative and fermentative pathways, we reasoned that the hypoxia-driven induction of fermentation might require a concomitant suppression of oxidation of the same substrates. To quantify BCAA oxidation, we measured the secretion of 3-hydroxyisobutyrate (3HIB), an intermediate in the valine oxidation pathway (**Figure 4A**).^37,49^ Consistent with prior work showing that hypoxic adipocytes suppress mitochondrial BCAA oxidation,^40^ we observed that hypoxic cells secreted approximately half the levels of valine-derived 3HIB compared to normoxic cells, which was further suppressed by Bcatm inhibition (**Figure 4B, S6A**).

**Figure 4.**
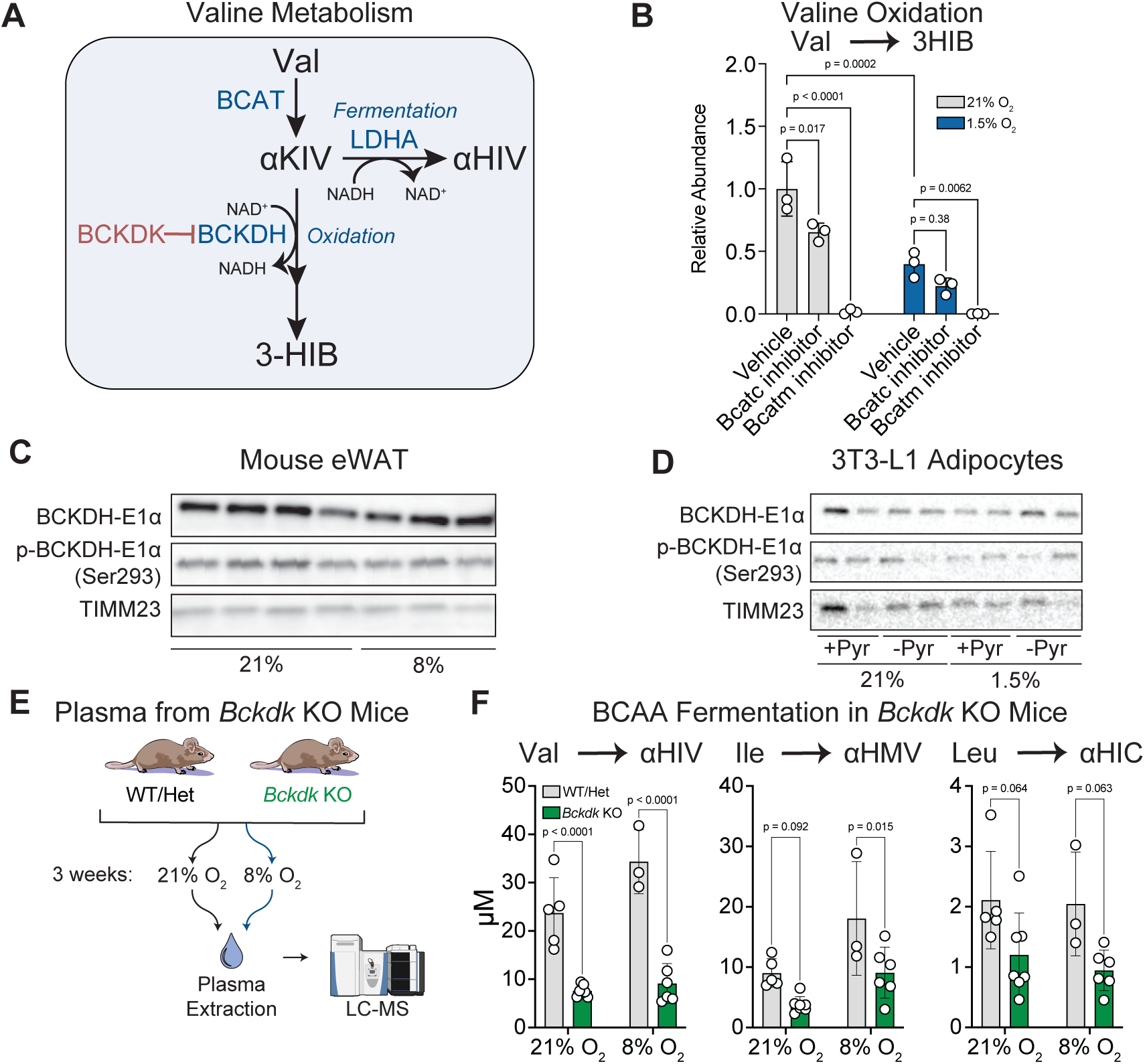
Restrained BCAA oxidation is required for BCAA fermentation *in vivo*. (A) Schematic depicting valine catabolism pathways, including regulation by Bckdk and the production of 3-HIB during valine oxidation. (B) Relative abundance of ^13^C-3HIB in conditioned media from 3T3-L1 adipocytes after 3-day incubation in 21% O_2_ or 1.5% O_2_ and treatment with 10 µM BCATc inhibitor 2 or 50 µM BCAT-IN-2 (BCATm inhibitor). n = 3 wells. (C-D) Immunoblots showing Bckdh-E1a, phospho-Bckdh-E1a (Ser 293), and Timm23 abundance in eWAT from mice housed at 21% O_2_ or 8% O_2_ for 24 hours (C) and in 3T3-L1 adipocytes cultured for 3 days in 21% O_2_ or 1.5% O_2_ with or without 1 mM sodium pyruvate (D). (E) Schematic depicting experiment for determining BCHA levels in wild-type and *Bckdk*-deficient mice. (F) Concentration of BCHAs in plasma from wild-type and *Bckdk*-deficient male mice housed at 21% O_2_ or 8% O_2_ for 2 weeks. WT/Het n = 5 mice, 21% KO n = 6 mice, 8% WT/Het n = 3 mice, 8% KO n = 6 mice. Mean ± SD are shown. Significance was determined by two-way ANOVA followed by Tukey’s multiple comparisons test (B) or Šídák’s multiple comparisons test (F).

BCKA oxidation is gated by the BCKDH complex, which catalyzes the irreversible commitment step (**Figure 4A**).^37^ BCKDH belongs to the mitochondrial α-ketoacid dehydrogenase family, which also includes pyruvate dehydrogenase (PDH). In hypoxia, PDH activity is suppressed by the upregulation of PDH kinases (PDKs) to divert pyruvate toward fermentation.^17^ The BCKDH complex is similarly regulated by a dedicated kinase, BCKDH kinase (BCKDK),^50^ which inhibits BCKDH activity and controls the rate of BCAA oxidation.^51,52^ We initially hypothesized that the shift in BCAA oxidation levels was mediated by the inhibitory phosphorylation of Bckdh (**Figure 4A**).^37^ However, contrary to the PDH paradigm, we observed no significant changes in total Bckdh levels or phosphorylation status in either mouse eWAT or in cultured adipocytes after exposure to hypoxia (**Figure 4C-D**). This finding suggests that the hypoxic suppression of oxidation is driven not by an increase in Bckdk activity, but by alternative regulatory mechanisms, such as direct inhibition by NADH.^50^

Nevertheless, based on the competition for the shared BCKA pool, we reasoned that activating BCAA oxidation by ablating the kinase Bckdk could suppress fermentation. To test this model, we exposed wild-type and *Bckdk*-deficient male mice^51^ to normoxia and hypoxia and measured plasma BCHA levels (**Figure 4E**). Consistent with the competition model, we observed that *Bckdk*-deficient mice displayed significantly lower circulating levels of BCHAs in both normoxia and hypoxia (**Figure 4F**). Therefore, we concluded that the Bckdk-imposed restraint on BCAA oxidation is required for the diversion of BCAAs into the fermentative pathway. Phenotyping of *Bckdk*-deficient mice revealed no overt deficits in their adaptation to hypoxia (**Figure S6B-G**).

### BCHAs are biomarkers of NADH reductive stress

Having established the mechanistic basis of BCAA fermentation, we investigated the physiological relevance of this pathway in humans by measuring BCHA levels in conditions characterized by acute NADH reductive stress. We focused our analysis on three distinct contexts known to induce reductive stress (**Figure 5A**):

1. resistance exercise, which involves short bursts of maximal effort and requires anaerobic energy production in skeletal muscle,
2. severe COVID-19, which induces systemic hypoxemia, and
3. alcohol consumption, which drives rapid hepatic NADH generation via alcohol dehydrogenase.^10,11^

**Figure 5.**
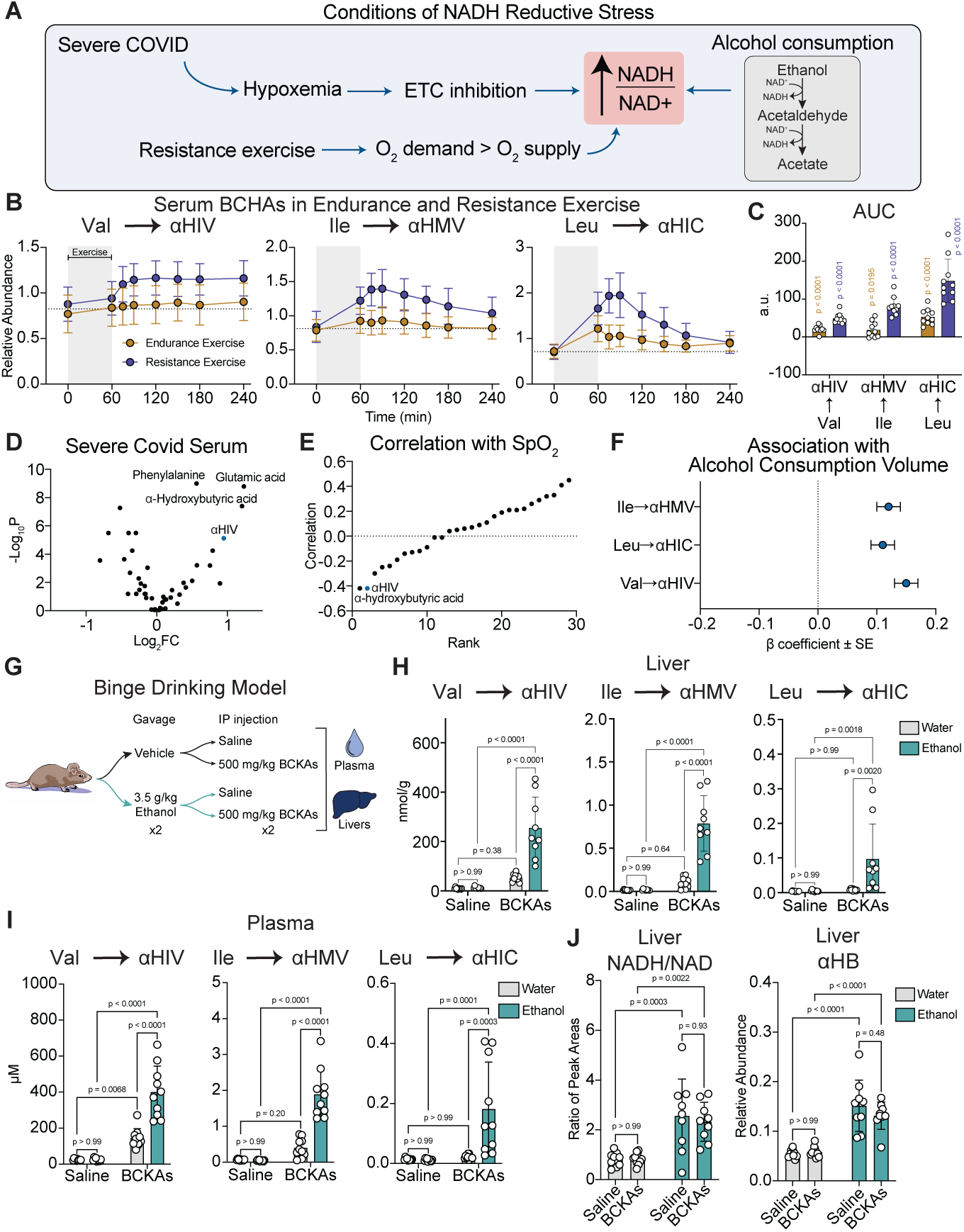
BCAA fermentation products are biomarkers of NADH reductive stress. (A) Schematic depicting the conditions inducing NADH reductive stress. (B) Relative abundance of BCHAs in serum of people undergoing endurance or resistance exercise immediately prior to their bout of exercise or up to 3 hours after (Morville et al., 2020). n = 10 human males. (C) Areas under the curve for circulating BCHA levels in people undergoing endurance or resistance exercise. (D) Enrichment of metabolites in serum from severe COVID patients compared to healthy controls (Páez-Franco et al., 2021). n = 27 control patients, n = 46 severe COVID patients. (E) Spearman correlation coefficient between serum levels of metabolites and peripheral O_2_ saturation. (F) â coefficient ± standard error in a linear model estimating the association between self-reported alcohol consumption and fasting serum levels of BCHAs among African American patients from the Jackson field center in the Atherosclerosis Risk in Communities Study (Zheng et al., 2014). n = 1500 patients. (G) Schematic depicting binge drinking model in mice. (H-I) Concentration of BCHAs in the liver (H) and plasma (I) of mice gavaged twice with either a vehicle control or 3.5 g/kg of ethanol and injected intraperitoneally twice with either saline or 500 mg/kg of each branched-chain ketoacid (BCKA): sodium α-ketoisovalerate, sodium α-ketoisocaproate, and sodium α-keto-β-methylvalerate. Water groups n = 10 mice, ethanol groups n = 9 mice. (J) NADH/NAD ratio and the relative abundance of α-hydroxybutyrate in mice gavaged with either a vehicle control or 3.5 g/kg of ethanol and injected intraperitoneally with either saline or 500 mg/kg of BCKAs. Water groups n = 10 mice, ethanol groups n = 9 mice. Mean ± SD are shown; mean ± SE are shown for F. Significance was determined using one-sample t-tests (c) and two-way ANOVA followed by Tukey’s multiple comparisons test (h-j). For metabolite measurements in COVID patients (d), significance was assessed using the Kruskal-Wallis test followed by Dunn’s post-hoc test (Páez-Franco et al., 2021).

To investigate the metabolic response to exercise, we analyzed published serum metabolite profiles from human subjects undergoing acute bouts of resistance or endurance exercise.^53^ Based on data from samples collected immediately prior to and over the 3 hours following physical activity, resistance exercise induced a significant increase in the circulating levels of all three BCHAs. In contrast, endurance exercise, which requires primarily aerobic oxidative metabolism, produced a much more muted increase in BCAA fermentation (**Figure 5B-C**).

We next examined the impact of pathological hypoxia by analyzing published serum metabolite profiles of patients suffering from severe respiratory distress due to COVID-19.^54^ Strikingly, αHIV (derived from valine) was identified as one of the most highly enriched metabolites in the serum of patients with severe symptoms (**Figure 5D**). Furthermore, αHIV levels exhibited a strong inverse correlation with blood oxygen saturation (SpO_2_) (**Figure 5E**), providing clinical evidence that valine fermentation scales quantitatively with the severity of systemic hypoxemia. Unfortunately, other BCHAs were not measured in this dataset.

Finally, we assessed the relationship between hepatic reductive stress and BCAA fermentation using data from a sample of African American subjects in the Atherosclerosis Risk in Communities (ARIC) study.^55^ In this dataset, circulating BCHA levels were significantly and positively associated with self-reported alcohol consumption volume (**Figure 5F**), consistent with the NADH burden imposed by hepatic ethanol oxidation.

To causally link ethanol oxidation to BCAA fermentation, we designed a combined binge drinking/BCKA challenge experiment in mice. Mice received two 3.5 g/kg doses of ethanol separated by one hour, with each dose followed by an intraperitoneal bolus of a mixture containing 500 mg/kg of each BCKA (**Figure 5G**). As expected, BCKA supplementation in mice not treated with ethanol caused a rapid accumulation of circulating BCKAs and a sustained elevation of BCHAs (**Figure S7A-B**).

In mice supplemented with BCKAs, ethanol intake increased the hepatic and plasma concentrations of BCHAs up to 12-fold (**Figure 5H-I**), confirming that binge drinking promotes BCAA fermentation. As expected, ethanol consumption also increased the hepatic NADH/NAD^+^ ratio and the levels of αHB, a marker of hepatic reductive stress (**Figure 5J**). However, BCKA supplementation did not alleviate this reductive stress. These results suggest that while BCAA fermentation is activated during hepatic reductive stress, the rate of BCHA production is likely too slow to offset the rapid NADH buildup from ethanol oxidation.

### Sperm-specific lactate dehydrogenase C drives BCAA fermentation

While our initial metabolomic profiling focused on male mice, we sought to determine if this metabolic adaptation was conserved across sexes. Surprisingly, female mice exhibited significantly lower circulating concentrations of BCHAs compared to males, with no significant increase in hypoxia (**Figure 6A**). To understand this sex-specific effect, we exposed castrated male mice to hypoxia and observed a marked reduction in circulating BCHAs compared to sham-operated controls (**Figure 6B**). Furthermore, quantification of absolute BCHA concentrations across tissues revealed that the testes contained the highest basal levels of BCHAs (**Figure 6C**), explaining the sexual dimorphism we observed. Testicular BCHA levels declined upon hypoxic exposure; viewed in combination with the castration data, this observation suggests that hypoxia triggers increased export of BCHAs from the testes into systemic circulation.

**Figure 6.**
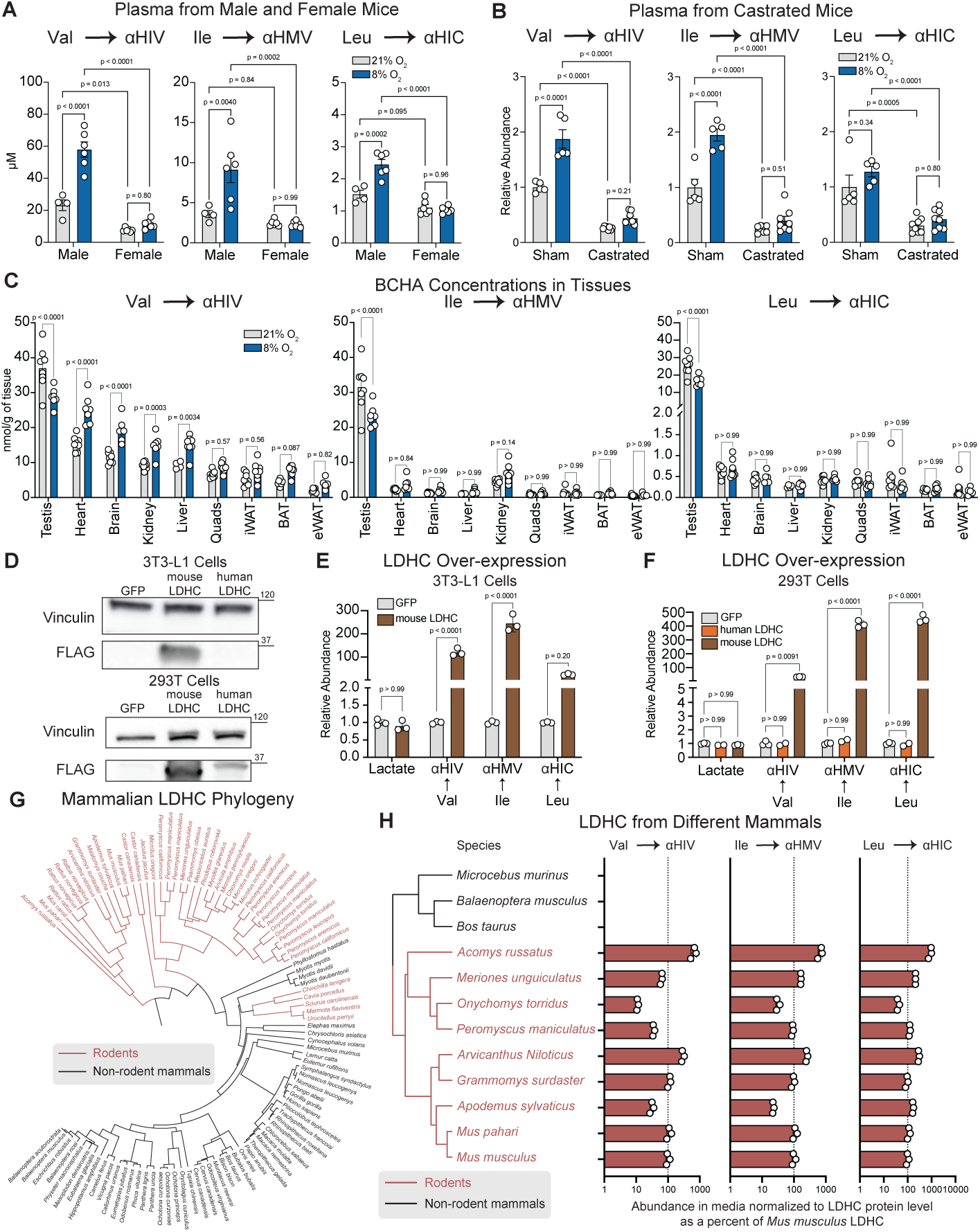
Sperm-specific lactate dehydrogenase C drives BCAA fermentation. (A) Concentration in plasma of BCHAs in male or female mice housed at 21% O_2_ or 8% O_2_ for 24 hours. 21% males n = 4; 21% females, 8% males and females, n = 6 mice. (B) Relative abundance in plasma of BCHAs from sham or castrated male mice housed at 21% O_2_ or 8% O_2_ for 24 hours. Sham n = 5 mice, castration n = 8 mice. (C) Concentration of BCHAs in tissues from male mice housed at 21% O_2_ or 8% O_2_ for 24 hours. In 21% O_2_, liver n = 4, every other organ n = 8. In 8% O_2_, eWAT n = 5; brain n = 6; testis, heart, kidney n = 7; liver, quads, iWAT, BAT n = 8. (D) Immunoblots of undifferentiated 3T3-L1 cells and HEK293T cells overexpressing GFP, mouse LDHC, or human LDHC. (E-F) Relative abundance of lactate and ^13^C-BCHAs in conditioned media from undifferentiated 3T3-L1 cells (E) overexpressing GFP or mouse LDHC and from HEK293T cells (F) overexpressing GFP, human LDHC, or mouse LDHC after 3 days of incubation at 21% O_2_. n = 3 wells. n = 2 wells for HEK293T cells over-expressing human LDHC. (G) Phylogenetic tree of LDHC from M*us musculus* and the 100 closest protein sequences. (H) Phylogenetic tree of 12 mammalian LDHC sequences and the relative abundance of ^13^C-BCHAs in the conditioned media from undifferentiated 3T3-L1 cells overexpressing each of the different LDHC sequences. Relative abundances were normalized based on LDHC protein expression, determined by immunoblot, and scaled to the metabolite abundance observed from the cells overexpressing *Mus musculus* LDHC. n = 3 wells. Mean ± SD are shown. Significance was determined by two-way ANOVA followed by Tukey’s multiple comparisons test (A, B, D, F) or Šídák’s multiple comparisons test (C, E).

Notably, the testis is the exclusive site of expression for lactate dehydrogenase C (LDHC) (**Figure S8A**).^56^ Older research has demonstrated that LDH from murine testes catalyzes the reduction of a broader array of substrates than LDHA or LDHB.^57^ To determine if LDHC possesses unique catalytic efficiency for BCAA fermentation, we attempted to overexpress murine *Ldhc* or human *LDHC* in 3T3-L1 and HEK293T cells but observed overexpression of human LDHC only in the 293T cells (**Figure 6D**). Strikingly, overexpression of murine LDHC dramatically increased BCAA fermentation in both cell lines, whereas over-expression of human LDHC failed to elicit a similar response (**Figure 6E-F, S8B-C**).

Given the sporadic distribution of BCAA fermentation activity across diverse kingdoms of life, including in fermentative bacteria and eukaryotic parasites,^58^ we asked whether this pathway evolved independently within the rodent lineage. To reconstruct the evolutionary history of this activity, we generated a phylogenetic tree utilizing the 100 protein sequences most homologous to *Mus musculus* LDHC. The resulting analysis revealed that almost all identified rodent LDHC orthologs clustered into three clades separated from the broader family of mammalian LDHC enzymes (**Figure 6G**).

Next, we asked whether the capacity for BCAA fermentation is a general property of rodent LDHC or restricted to mice. We expressed a library of 12 LDHC orthologs—9 from rodents and 3 from non-rodent mammals—in 3T3-L1 cells (**Figure S9A**). Every tested rodent LDHC construct conferred robust BCAA fermentation, whereas orthologs from non-rodent mammals (including blue whale, mouse lemur, and cattle) failed to catalyze detectable BCHA production (**Figure 6H**, **S9B**).

Based on these findings, we conclude that the capacity to efficiently ferment BCAAs evolved convergently across diverse organisms, including certain fermentative bacteria,^34–36,59^ halophilic archaea,^60^ unicellular eukaryotes,^61,62^ and rodent sperm (**Figure S9C**). Notably, BCAA fermentation has been characterized in the flagellated protozoan parasites *Trypanosoma cruzi* and *Trichomonas vaginalis*. In *T. cruzi*, pharmacological inhibition of this pathway compromises motility and viability.^63,64^ Intriguingly, *Ldhc*-deficient mouse sperm also exhibit impaired motility,^65^ suggesting that BCAA fermentation may play a similarly important role in supporting rodent sperm bioenergetics and motility in the female reproductive tract.

### Branched-chain hydroxyacids are anaerobic electron carriers facilitating hyperactive motility in mouse sperm

Having demonstrated that rodent LDHC facilitates efficient BCAA fermentation, we asked whether this pathway functions as a physiologically significant anaerobic electron sink in mouse sperm. To answer this question, we treated isolated mouse sperm with the Complex I inhibitor rotenone or the Complex IV inhibitor cyanide as models of reductive stress (**Figure 7A**). In each condition, sperm were supplemented with either pyruvate, a known electron acceptor, or a mixture of BCKAs. As expected, ETC inhibition elevated the intracellular NADH/NAD^+^ ratio by up to 3-fold. However, supplementation with either pyruvate or BCKAs significantly lowered this ratio, rendering the sperm resistant to the redox perturbations induced by respiratory blockade (**Figures 7B-C**).

**Figure 7.**
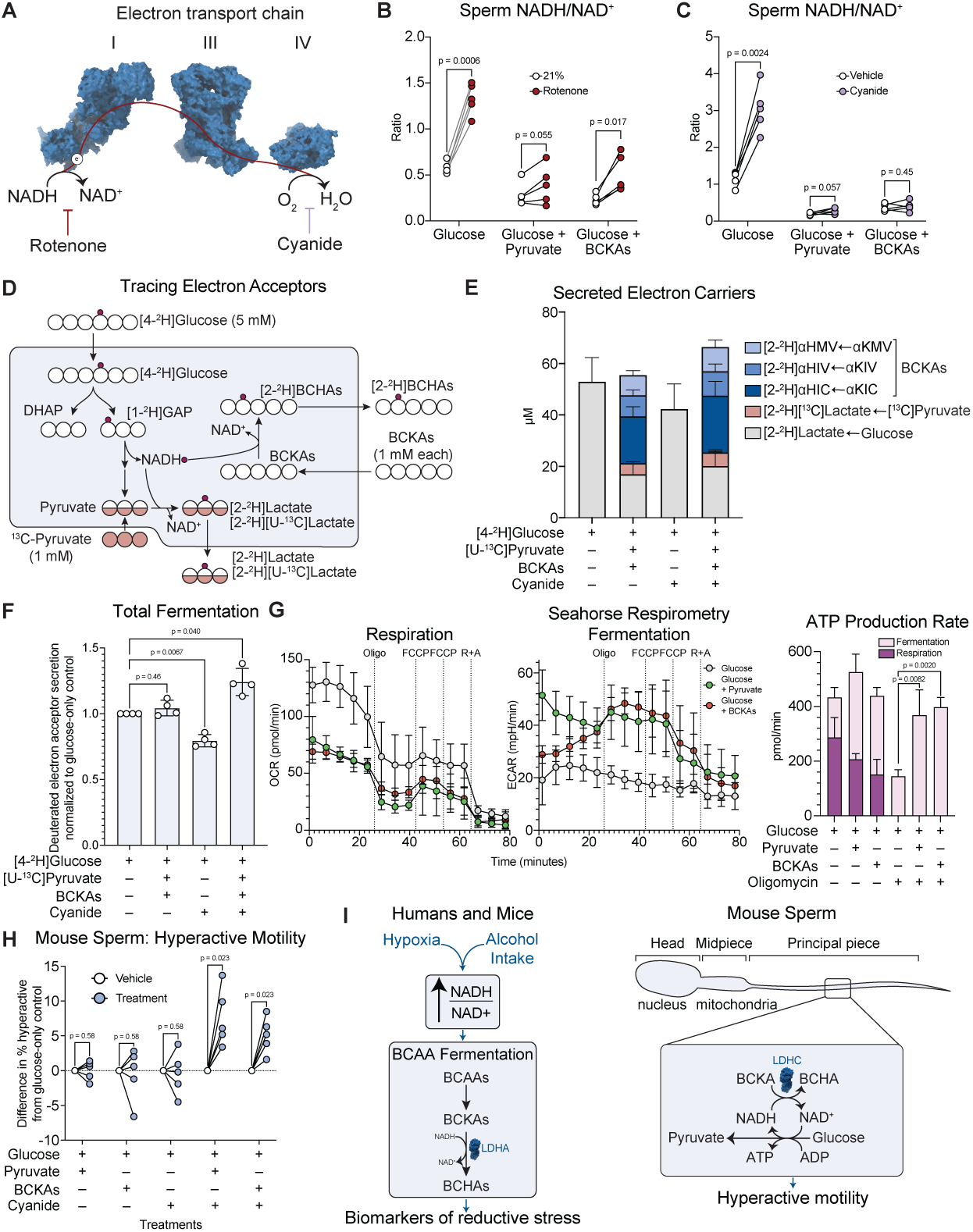
Branched-chain hydroxyacids are anaerobic electron carriers facilitating hyperactive motility in mouse sperm. (A) Schematic depicting inhibition of complexes I and IV of the electron transport chain. (B-C) NADH/NAD ratio in capacitated mouse sperm cultured with 5.54 mM glucose, supplemented with either 5 mM pyruvate or 1.67 mM of each BCKA. Sperm were treated with either 0.1% ethanol or 500 nM rotenone (B) or water or 10 mM sodium cyanide (C). n = 5 mice. Equal numbers of sperm from each mouse were split into each condition. (D) Schematic depicting [4-^2^H]glucose tracer experiment in mouse sperm supplemented with [U-^13^C]pyruvate and BCKAs. (E) Concentration of ^2^H-labeled hydroxyacids in the media from capacitated mouse sperm cultured with 5 mM [4-^2^H]glucose with or without 1 mM of [U-^13^C]pyruvate and 1 mM of each BCKA. Sperm were cultured for 3 hours and were supplemented with either a vehicle control or 10 mM sodium cyanide. n = 4 mice. Equal numbers of sperm from each mouse were split into each condition. (F) Relative fermentation levels of capacitated sperm cultured with 5 mM [4-^2^H]glucose with or without 1 mM of [U-^13^C]pyruvate and 1 mM of each BCKA. Relative fermentation level was calculated as the total concentration of ^2^H-labeled hydroxyacids in the media, scaled to the condition in which sperm were cultured with [4-^2^H]glucose as the only carbon source. n = 4 mice. Equal numbers of sperm from each mouse were split into each condition. (G) Oxygen consumption rate, extracellular acidification rate, and inferred ATP production rates of non-capacitated mouse sperm cultured with 5 mM glucose supplemented with either 5 mM pyruvate or 1.67 mM of each BCKA. Sperm were treated with the complex V inhibitor oligomycin (5 µM), two doses of the uncoupler FCCP (600 nM), and a combined dose of the complex I inhibitor rotenone (1 µM) and complex III inhibitor antimycin A (1 µM). n = 4 wells. Sperm from 2 mice were pooled and split into each well. (H) Difference in % hyperactive out of all detected motile trajectories comparing each treatment condition to glucose-only control. n = 5 mice. Equal numbers of sperm from each mouse were split into each condition for capacitation. (I) Schematic showing the role of BCAA fermentation as a biomarker of reductive stress in humans and mice (left) and as an electron sink facilitating hyperactive motility in the principal piece of mouse sperm (right). Mean ± SD are shown. Significance was determined using paired t-tests corrected for multiple comparisons using False Discovery Rate (B, C, H) or repeated-measures one-way ANOVA followed by Dunnett’s multiple comparisons test (F) or two-way ANOVA followed by Tukey’s multiple comparisons test (G).

To orthogonally validate that BCKAs directly consume cytosolic reducing equivalents, we performed a [4-^2^H]glucose tracer experiment in mouse sperm. During glycolysis, the oxidation of [4-^2^H]glucose generates cytosolic NADH carrying the ^2^H label. Consequently, any metabolite that accepts electrons from cytosolic NADH will incorporate this ^2^H label.^66,67^ We cultured sperm with [4-^2^H]glucose alone or supplemented with unlabeled BCKAs and [U-^13^C]pyruvate (labeled to distinguish exogenous uptake from glycolytic production) (**Figure 7D**).

In the absence of exogenous electron acceptors, [2-^2^H]lactate generated from anaerobic glycolysis was the sole secreted electron carrier (**Figure 7E**). The addition of pyruvate and BCKAs significantly diverted reducing equivalents away from glycolytic lactate and resulted in the secretion of [2-^2^H]BCHAs and [2-^2^H][U-^13^C]lactate. In this condition, labeled BCHAs comprised more than half of the secreted electron carriers and were secreted at similar concentrations to lactate, indicating that BCAA fermentation serves as a primary electron sink even in the presence of other electron acceptors. When respiration was inhibited, the total secretion of labeled fermentation products increased only in conditions where pyruvate and BCHAs were supplied (**Figure 7F**). Taken together, these results indicate that sperm use BCHA production as a high-flux electron sink to increase fermentation during respiratory blockade.

To quantify real-time respiration and fermentation rates in each condition, we performed respirometry on isolated, attached sperm using a Seahorse extracellular flux analyzer. Supplementation with either pyruvate or BCKAs suppressed basal respiration by approximately 40% (**Figure 7G**), a finding confirmed in freely moving sperm using high-resolution Oroboros respirometry (**Figure S10A-B**). This respiratory suppression is consistent with a model where high-affinity fermentative dehydrogenases divert electrons away from the ETC. In turn, fermentation significantly increased following oligomycin (Complex V inhibitor) treatment *only* when sperm were supplemented with either pyruvate or BCKAs (**Figure 7G**). These data suggest that by providing an immediate sink for electrons, exogenous BCKAs increase the maximal fermentation rate of mouse sperm.

Finally, we investigated the physiological consequences of this metabolic flexibility on sperm motility. As sperm traverse the female reproductive tract, they undergo capacitation, a biochemical maturation process that induces hyperactive motility, characterized by non-linear, whiplash movement of the flagellum. Hyperactive motility is required for passing through the zona pellucida, the dense matrix surrounding the egg.^68^ We assessed hyperactive motility across paired conditions manipulating respiration and fermentation. Strikingly, cyanide treatment or supplementation with pyruvate or BCKAs had little effect on hyperactive motility. However, maximizing fermentation by inhibiting respiration and supplying electron acceptors (either pyruvate or BCKAs) significantly enhanced hyperactive motility (**Figure 7H**, **S10E**). This fermentative state may represent the bioenergetic conditions of capacitated sperm in the oviduct.

These results support the model that fermentation is a privileged power source for hyperactivation. Indeed, prior work has demonstrated that promoting glycolysis increases hyperactive motility^69^ and that impaired glycolysis suppresses fertilization by murine sperm.^70,71^ Interestingly, respiration and fermentation are highly compartmentalized within spermatozoon. Glycolytic enzymes driving fermentation are tethered to the fibrous sheath of the flagellar principal piece,^72,73^ placing them in direct proximity to the dynein motors powering whiplash beating of the flagellum. In contrast, mitochondrially derived ATP is restricted to the midpiece and diffuses too slowly to reach distal motors.^74,75^ Consistent with this hypothesis, murine sperm depleted of mitochondrial ATP retain hyperactive motility.^76^

In summary, our findings establish BCAA fermentation as a conserved biomarker of cytosolic reductive stress. While this pathway acts as a latent sink for excess electrons in most tissues, its high-flux activation in rodent sperm represents an evolutionary innovation to support anaerobic energy production, which is a universal metabolic requirement for hypermotility across mammalian sperm (**Figure 7I**).

## DISCUSSION

In this study, we develop a rich resource uncovering the tissue-specific metabolomic features of mammalian hypoxia adaptation. Using this resource, we identify a fermentative branch of BCAA catabolism that is robustly activated in hypoxia and other conditions of cytosolic reductive stress. In most mammalian organs, this pathway relies on the catalytic promiscuity of lactate dehydrogenase A, serving as an exquisitely sensitive biomarker of reductive stress. In rodent sperm, lactate dehydrogenase C has evolved to catalyze BCAA fermentation with much higher efficiency. This enables BCAA fermentation as a physiologically significant anaerobic electron sink in the sperm flagellum, providing localized NAD^+^ regeneration to support hyperactive motility.

Although BCHAs are structurally similar to lactate, they fulfill a unique physiological role. Because LDHA exhibits a relatively high K_m_ for BCKAs, the rate of BCHA production is low at baseline. This feature makes the pathway highly responsive to substrate availability. As the intracellular NADH/NAD^+^ ratio rises, BCHAs accumulate, establishing these metabolites as sensitive biomarkers for cytosolic reductive stress. Consistent with this model, BCHA levels exhibited a greater fold-change enrichment in hypoxia than most other electron acceptors and were significantly elevated across diverse models of reductive stress in humans, including resistance exercise, severe COVID-19, and alcohol consumption.

Unlike lactate, which is readily consumed as a major circulating fuel source,^30,31^ BCHAs primarily represent an excretable waste product. Whereas lactate was undetectable in urine, BCHAs were found at similar or greater levels in urine than in plasma. Therefore, the excretion of BCHAs provides a mechanism to permanently eliminate excess electrons from the organism. A similar strategy for disposing of electron carriers has been observed in the anoxia-tolerant goldfish, which excretes ethanol derived from fermentation.^21^

While amino acid fermentation remains under-appreciated in mammalian metabolism, the use of amino acid-derived metabolites as electron acceptors is an ancient biological strategy. In anoxic oyster hearts, for example, aspartate is fermented following a similar strategy to BCAA fermentation. The de-amination of aspartate yields oxaloacetate, which accepts electrons from cytosolic NADH to form malate and allow continuous glycolytic ATP production.^24,25^ Similarly, obligate anaerobic bacteria belonging to the genus *Clostridium* rely almost exclusively on amino acids for energy production. In a process termed Stickland fermentation, these organisms couple the oxidation of one amino acid to the reduction of another, maintaining redox balance while driving substrate-level phosphorylation.^77–79^ Notably, the reductive metabolism of leucine in this context involves the production of a BCHA as an electron sink, directly mirroring the fermentative strategy observed in hypoxic mammals.^36^ These pathways have been proposed as some of the earliest energy-producing processes, given that amino acids were likely more abundant than carbohydrates on the primitive, hypoxic planet.^79,80^

In modern mammals, fermentative capacity is largely constrained by the substrate specificity of lactate dehydrogenase enzymes, which typically favor pyruvate. However, the rodent LDHC evolved to relax substrate specificity and efficiently reduce a broader set of keto-acids, including BCKAs. This specialization may have been driven by the intense sperm competition unique to rodents, which selects for velocity over longevity. To achieve higher velocities, rodent sperm evolved longer flagella,^74,81^ pushing distal motor proteins farther from the mitochondria-rich midpiece. In this context, a high-affinity fermentation pathway capable of utilizing a broader set of electron acceptors may have supplied a bioenergetic advantage. The importance of these alternative electron acceptors for hypermotility may partially explain the observed sub-fertility in male *Bckdk*-knockout rats^82^ and mice,^83^ which exhibit decreased BCAA fermentation.

The convergent evolution of BCAA fermentation in phylogenetically diverse flagellated eukaryotes exemplifies how bioenergetic pressure imposed by specialized functions (e.g. flagellar motility) drives metabolic convergence. Our findings underscore that ancient metabolic pathways can be resurrected by evolution to support anaerobic metabolism. We anticipate that the time- and organ-resolved metabolomics resource published here will help uncover other metabolic adaptations underlying the mammalian response to hypoxia. A more comprehensive understanding of these anaerobic pathways will reveal new opportunities in the diagnosis and treatment of diseases caused by hypoxia and other conditions of reductive stress.

## RESOURCE AVAILABILITY

### Lead Contact

Further requests and information concerning this study should be addressed to the lead contact, Isha H. Jain.

### Materials availability

All the reagents are available upon request to the lead contact, Isha H. Jain.

### Data and code availability

Metabolomics data are available at https://jain-lab-ucsf.github.io/hypoxia-metabolomics/. Other data are available upon request.

## ACKNOWLEDGEMENTS

We thank all members of the Jain lab for thoughtful discussions and review of the manuscript. We thank Yuyin Zhou for her guidance with mass spectrometry and her advice during project conceptualization. We thank Shweta Bhagwat and Celia Santi for their assistance with mouse sperm motility quantification. We thank the Innovation Core at the Weill Institute for Neuroscience for their assistance with imaging. We thank Valentin Cracan for sharing an *Lb*NOX over-expression plasmid. We acknowledge Hanifa Mohammed for assistance with generating and validating knockout lines. We thank Brian Plosky and April Pawluk for their thoughtful insights during the writing process and Tami Tolpa for her assistance with graphic design.

Funding was provided by the National Institute of General Medical Sciences Medical Scientist Training Program, grant T32GM141323 (A.D.M. and A.P.), the National Institute of Diabetes, Digestive, and Kidney Diseases under grants 5F30DK139713 (A.D.M.) and 5R01DK109714 (T.G.A.), the National Science Foundation Graduate Research Fellowship program under grant no. 2034836 (B.T.L.C.), the National Heart, Lung, and Blood Institute under grants F31HL172581 (W.R.F.) and R01HL179702 (I.H.J.), National Institute on Aging R01AG065428 (W.R.F.), the Hillblom Foundation Award GR-02469 (I.H.J.), the Defense Advanced Research Projects Agency, Biological Technologies Office Program: Panacea issued by DARPA/CMO under Cooperative Agreement No. HR0011-19-2-0018 (I.H.J., B.B.Q., A.M.B., S.J.A., L.F.W.), and the National Institute of Neurological Disorders and Stroke 1F31NS139663 (A.P.). I.H.J. is a Core Investigator at Arc Institute, and her research is supported by Arc. Any opinions, findings, and conclusions or recommendations expressed in this material are those of the authors and do not necessarily reflect the views of the National Science Foundation or National Institutes of Health.

## AUTHOR CONTRIBUTIONS

A.D.M. and I.H.J. conceived the project and acquired funding for the project. A.D.M., B.T.L.C., Y.M.M., and S.Y.B. performed experiments with *Bckdk*-deficient mice and mice treated with ethanol. A.D.M., B.T.L.C., W.R.F., and A.G.H. performed genetic manipulations and isotope-tracing experiments *in vitro*. A.D.M., A.P., and P.L. conducted experiments involving primary mouse sperm. A.D.M., B.R.D., J.S., and M.K. performed respirometry experiments with primary mouse sperm. B.B.Q. and A.M.B. conducted experiments for hypoxia and FG-4592 metabolomics. T.G.A. provided the *Bckdk*-deficient mice. M.T. and R.T. performed data analysis of the untargeted metabolomics data. I.H.J., P.L., M.P., Y.K., S.J.A., and L.F.W. supervised study design, experimentation, and data analysis. A.D.M. and I.H.J. wrote the manuscript with input from all authors.

## DECLARATION OF INTERESTS

The authors declare no competing interests.

## METHODS

### Experimental model and study participant details

#### Animal studies

All animal studies were approved by the UCSF/Gladstone Institutes IACUC, and by the WashU Medicine Animal Care and Use Committee (protocol 22-0251). Most mouse experiments were performed at UCSF/Gladstone Institutes, where mice were housed at ∼20 °C with 50% humidity and a 12 hour light/dark cycle. Mice were fed a chow diet (PicoLab 5058). Wild-type C57BL/6J mice were purchased from the Jackson Laboratory.

For sperm motility experiments, 3-6-month-old male C57BL/6J mice were purchased from Charles River and housed at a WashU barrier animal facility in a room with a controlled dark-light cycle (10 hours dark followed by 14 hours of light) and a controlled temperature of approximately 23 ± 0.5°C. The animals were fed a standard chow diet and provided with chlorinated water *ad libitum*.

Mice were euthanized by asphyxiation with CO_2_ followed by cervical dislocation with every effort made to minimize animal suffering.

#### *In vivo* hypoxia treatment

Hypoxic environments were established in 472 L acrylic chambers by blending nitrogen produced by a generator (N2Gen-02CPi-P, South-Tek Systems) with compressed air to achieve oxygen concentrations of either 8% or 11% using a gas-mixing system. Humidity within the chambers was maintained using open water reservoirs.

In both conditions, inspired oxygen and carbon dioxide concentrations were continuously monitored with a wireless system and confirmed daily. Carbon dioxide accumulation was mitigated by placing soda lime (Fisher Scientific) inside each chamber. Animals were housed in their home cages, which were changed at least weekly, and had unrestricted access to food and water.

For untargeted metabolomics experiments, mice that were housed at 8% O_2_ were acclimated at 11% O_2_ for a week prior to the initiation of 8% O_2_ treatment. To induce HIF stabilization acutely, mice were treated with 100 mg/kg doses of FG-4592 12 hours apart. After the acute timepoints, 50 mg/kg doses of FG-4592 were administered 3 times weekly for up to 3 weeks.

#### Isolation and capacitation of mouse sperm

Capacitation solution was prepared with the following components (g/L): fatty acid-free fraction V bovine serum albumin (BSA) (10), CaCl_2_·2H_2_O (0.3), KCl (0.3496), KH_2_PO_4_ (0.0504), MgSO_4_·7H_2_O (0.0493), NaCl (5.9375), NaHCO_3_ (2.1), and HEPES (4.766). The solution was adjusted to pH 7.4 with NaOH; 320 ± 5 mOsm/L. This solution was supplemented with various nutrients as described for specific experimental methods.

Cauda epididymides were dissected from euthanized adult male mice and immediately placed into 2 mL of prewarmed capacitation medium. Multiple incisions were made in caudae epididymides followed by incubation at 37 °C in a humidified atmosphere containing 5% CO_2_ for 10 min to allow sperm to disperse. For capacitation, sperm were subsequently incubated at 37 °C and 5% CO_2_ for 90 min prior to downstream analyses.

#### Cell culture

3T3-L1 fibroblasts were maintained in complete media: DMEM (Thermo Fisher Scientific, 11995073) with 10% FBS (Fisher Scientific, MT35015CV) and 1% penicillin-streptomycin-glutamine (Fisher Scientific, MT30009CI). Fibroblasts were differentiated into adipocytes using the following protocol: 20,000 cells were plated per well in 24-well plates. After 2 days, the media was replaced with complete media containing 115 mg/L IBMX, 1 mg/L insulin, and 100 μM dexamethasone. After 2 more days, the media was replaced with complete media containing 1 mg/L insulin. Over the following 6 days, media was replaced with complete media every 2 days. At this stage, cells were treated with drugs or different oxygen levels.

*In vitro* hypoxia experiments were performed in a dedicated 1.5% O_2_ chamber. Nitrogen gas was supplied by a nitrogen generator (N2Gen-02CPi-P, South-Tek Systems). For HIF activation experiments *in vitro*, cells were treated with 75 μM FG-4592 or 0.15% DMSO. For complex I inhibition, cells were treated with 500 nM rotenone or 0.1% ethanol. For GOT inhibition, cells were treated with 400 µM AOA. For BCAT inhibition, cells were treated with 10 µM BCATc inhibitor 2, 50 µM BCAT-IN-2, or 0.1% DMSO. For complex IV inhibition, cells were treated with 10 mM cyanide.

### Method details

#### Sample preparation for untargeted metabolomics

Preparation of flash-frozen tissue samples was conducted by Metabolon. Automated sample preparation was conducted on a MicroLab STAR® platform (Hamilton Company) beginning with the addition of recovery standards. Metabolites were extracted and proteins precipitated via methanol addition and mechanical shaking (GenoGrinder 2000, 2 min), followed by centrifugation. The extract was partitioned into five fractions: three for RP/UPLC-MS/MS (two positive, one negative ESI), one for HILIC/UPLC-MS/MS (negative ESI), and one backup.

Following organic solvent removal via TurboVap® (Zymark), extracts were stored overnight under nitrogen before LC-MS analysis.

#### Mass spectrometry for untargeted metabolomics

Sample analysis for untargeted metabolomics was performed by Metabolon. Chromatographic analysis was performed using a Waters ACQUITY UPLC coupled to a Thermo Scientific Q-Exactive high-resolution/accurate mass spectrometer equipped with a HESI-II source and an Orbitrap analyzer (35,000 resolution). Dried extracts were reconstituted in method-specific solvents containing fixed concentrations of internal standards to monitor injection and chromatographic consistency. Four distinct methods were employed: (1) an acidic positive ion condition optimized for hydrophilic compounds using a C18 column (Waters BEH C18, 2.1×100 mm, 1.7 µm) eluted with water/methanol containing 0.05% PFPA and 0.1% FA; (2) an acidic positive ion condition optimized for hydrophobic compounds using the same C18 column eluted with a higher organic gradient of methanol/acetonitrile/water containing 0.05% PFPA and 0.01% FA; (3) a basic negative ion condition using a dedicated C18 column eluted with methanol/water and 6.5 mM ammonium bicarbonate (pH 8); and (4) a negative ionization HILIC method (Waters BEH Amide, 2.1×150 mm, 1.7 µm) using a water/acetonitrile gradient with 10 mM ammonium formate (pH 10.8). The MS analysis (70–1000 m/z) alternated between MS and data-dependent MS^n^ scans using dynamic exclusion.

#### Compound identification and quantification for untargeted metabolomics

Raw data extraction, peak identification, and quality control were performed by Metabolon. Metabolites were identified by comparison to a library of over 3,300 commercially available purified standards and recurrent unknown entities. Identification was based on three strict criteria: (1) retention index (RI) within a narrow window of the proposed identification, (2) accurate mass match to the library within ± 10 ppm, and (3) MS/MS forward and reverse scores based on spectral comparison between experimental data and authentic standards. Recurrent unnamed biochemicals were tracked and identified by their unique chromatographic and mass spectral signatures to ensure consistency across studies.

Metabolite peaks were quantified using area-under-the-curve (AUC) integration. For studies requiring multiple days of analysis, a "block correction" normalization was employed to account for inter-day instrument variability. This process involved registering the median of each compound to 1.00 for each run-day block and normalizing individual data points proportionately.

To calculate the mass-normalized and log-transformed data, a multi-step scaling and imputation pipeline was employed:

1. Batch Normalization: To account for inter-day instrument variability, raw metabolite values were first divided by the median value of their respective instrument batches, resulting in a batch-specific median of one.
2. Mass Normalization: These batch-normalized values were then divided by the specific sample mass to adjust for variations in the amount of starting material.
3. Median Re-scaling: Following mass correction, each metabolite was re-scaled to a median of one by dividing the values by the overall metabolite-specific median.
4. Imputation: Missing values were imputed using the minimum observed value for each specific metabolite across all samples. These values were considered “relative abundances” for the purpose of Figures 1B, C, H.
5. Log Transformation: Finally, as metabolomic data typically follow a log-normal distribution, the mass-normalized and imputed values were natural log-transformed. These “log-transformed abundance” values were used in Figure S2A.

Liver 11% and 8% O_2_ 1-week samples were collected, processed, and analyzed in a separate experiment alongside 21% O_2_ control samples and were median-normalized independently from all the other samples.

#### Analysis of untargeted metabolomics data

For each organ, we performed a principal component analysis (PCA) on the most variable metabolites (variance > 0.5) after centering each metabolite to its mean across all samples. Analysis used prcomp() in R; visualization used fviz_pca_ind() from factoextra (**Figure S1A**).

For each metabolite, a linear model tested the association between metabolite levels and timepoint (baseline, 3h, 24h, 1w, 3w) under 8% O_2_ or 11% O_2_ exposure. Model significance was assessed by ANOVA comparison to an intercept-only model, with FDR correction applied.

Metabolites with FDR ≤ 0.05 were considered significantly associated with timepoint; their linear model coefficients (effect sizes) were used for downstream analyses. Linear model coefficients for metabolites significant in ≥70% of organs were compiled across all organ-timepoint combinations (excluding 21% O_2_ controls) and analyzed by PCA (Figure S1B). Pairwise Pearson correlations between organs were calculated from the linear model coefficients of shared significant metabolites using rcorr() from Hmisc, visualized with corrplot (Figure S1C).

For each organ, metabolites with p ≤ 0.05 for the 8% O_2_ 24h coefficient (vs. 21% O_2_ 24h) were identified and counted by the number of organs in which they reached significance. We visualized the counts using a stacked barplot function implemented in ggplot2 (Figure S1D).

Metabolites significantly altered (FDR ≤ 0.05) by FG-4592 or 8% O_2_ 24h (each vs. 21% O_2_ 24h) were compared using Venn diagrams generated with ggVennDiagram (Figure S1E). Linear model coefficients for significant plasma metabolites were clustered using k-means (k = 6) and visualized as a heatmap using pheatmap. Cluster-specific kinetics were displayed as box plots using ggplot2 (Figure S1F).

#### ^13^C-BCAA stable isotope tracer experiments

DMEM powder without glucose, glutamine, isoleucine, leucine, valine, sodium pyruvate, sodium bicarbonate, phenol red (US Biologicals, D9800-36) was used as a base and supplemented with 25 mM glucose, 4.0 mM glutamine, 0.8 mM ^13^C-isoleucine (Cambridge Isotope Laboratories, 2248), 0.8 mM ^13^C-leucine (Cambridge Isotope Laboratories, 2262), 0.8 mM ^13^C-valine (Millipore Sigma, 758159), 44 mM sodium bicarbonate, 10 mM HEPES, and 0.04 mM phenol red.

Complete media was prepared by adding dialyzed FBS (Thermo Fisher, 26400044) and penicillin-streptomycin-glutamine (Fisher Scientific, MT30009CI) to a final concentration of 10% and 1% respectively.

After 24 hours of treatment (with hypoxia or relevant drugs) in unlabeled media, the media was replaced with labeled media. Media samples were collected at 24, 48, or 72 hours following the isotope switch.

#### Targeted measurement of BCHAs and lactate

Polar metabolites were extracted from media, plasma, tissue, and standards by mixing 20 μL of liquid samples or standards with 200 μL of 80% methanol with 1 mM of the internal standard

D8-Valine. Plasma was extracted from blood by centrifuging at 2,000 rpm for 10 minutes at 4 °C and collecting the supernatant. Flash-frozen liver samples were pulverized using a cryo-cup grinder, and 20 mg of each sample was added to 200 μL of 80% methanol with 1 mM D8-

Valine. For each sample type, the mixture was incubated at −80 °C for 3 hours and vortexed. Next, the mixtures were centrifuged for 10 minutes at 16,000 × g at 4 °C. Supernatants were transferred to glass vials for storage or analysis.

Samples were analyzed on an Orbitrap Exploris 240 high-resolution mass spectrometer (Thermo Fisher Scientific) equipped with an electrospray ionization source and coupled to a Vanquish Horizon UHPLC system. Polar metabolites (5 μL per sample) were separated using reverse phase chromatography on a Hypersil GOLD C18 column (2.1 × 150 mm, 1.9 µm; Thermo Scientific 25002-152130) connected to a Hypersil GOLD C18 guard column (2.1 x 10 mm, 3 µm; Thermo Scientific 25003-012101). The autosampler was maintained at 4 °C and the column temperature at 45 °C. Mobile phase A consisted of water with 0.1% formic acid, while mobile phase B consisted of methanol with 0.1% formic acid. Separation was performed at a flow rate of 300 μL/min using the following gradient: 0 to 50% B from 0-8 min, 50% to 98% B from 8-9 min, hold at 98% B from 9-13 min, 98 to 0% B from 13-13.1 min, and hold at 0% B from 13.1-15 min. Each run was preceded by a 5-min equilibration at initial conditions.

The mass spectrometer was operated in negative mode using a spray voltage of 3 kV with sheath gas at 40, aux gas at 8, sweep gas at 1, ion transfer tube at 275 °C, and vaporizer at 320 °C. The mass spectrometer was operated in either full scan mode or targeted selected ion monitoring (tSIM) mode. In full scan mode, the resolution was 60,000, scan range was 70-100 m/z, RF lens was 60%, the AGC target was 1e7, and maximum injection time was 100 ms. In tSIM mode, the isolation window was 0.4, the resolution was 60,000, RF lens was 60%, AGC target was 1e6, and maximum injection time was 30 ms. In tSIM mode, the following compounds were detected (with retention time range in minutes): D8-valine (0-3), ^13^C-αHIV (5-9), ^13^C-αHMV/^13^C-αHIC (7-11), and ^13^C-3HIB (2.5-5.5). Relative abundances were calculated by dividing the peak area of the compound of interest by the peak area of D8-valine from the same sample and scaling to the control condition.

#### Genetic deletions

Guide RNA sequences were selected using CRISPick.^84,85^ Two guides were used for each knockout:

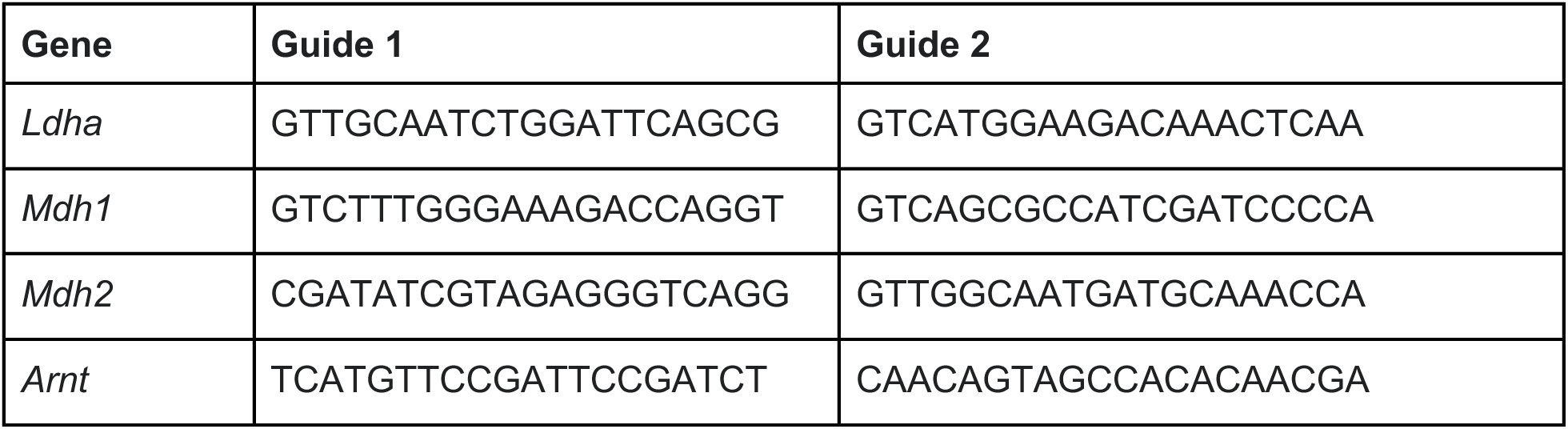

We digested the lentiCRISPRv2 backbone (Addgene 52961) ligated with the appropriate guide RNA sequences. Recombinant plasmids were packaged into lentiviral particles by co-transfection into HEK293T cells together with the helper plasmids pVSV-G (Addgene 8454) and psPAX2 (Addgene 12260). Viral supernatants were used to transduce 3T3-L1 cells via spinfection. Transduced cells were selected with puromycin (8 µg/mL for 3-5 days). Polyclonal populations of infected cells were used for downstream experiments.

#### Genetic over-expression

Mouse LDHA, human LDHA, and *Weissella* L-hydroxyisocaproate dehydrogenase proteins were over-expressed using EF1α-puromycin lentiviral vectors (Twist Bioscience). *Lb*NOX and LDHC plasmids were over-expressed using CMV-blasticidin lentiviral vectors (Twist Bioscience). Recombinant plasmids were packaged into lentiviral particles by co-transfection into HEK293T cells together with the helper plasmids pVSV-G (Addgene 8454) and psPAX2 (Addgene 12260). Viral supernatants were used to transduce 3T3-L1 cells via spinfection.

Transduced cells were selected with puromycin (8 µg/mL for 3-5 days) or blasticidin (10 µg/mL for 3-5 days).

#### Human genetics data analysis

Metabolite GWAS data from the METSIM study^42^ were processed from the PheWeb portal: https://pheweb.org/metsim-metab/. eQTL data were analyzed using the FIVEx browser (https://fivex.sph.umich.edu/), which includes data from 16 different studies.^44^

#### Immunoblotting

Cells were lysed in ice-cold RIPA buffer (Thermo Fisher Scientific) containing Halt Protease and Phosphatase Inhibitor Cocktail. Protein concentrations were measured using a Pierce BCA Protein Assay Kit (Thermo Fisher Scientific) and normalized across samples. Lysates were mixed with 6× Laemmli SDS sample buffer (Thermo Fisher Scientific) and heated at 95 °C for 5 min. Equal amounts of protein were resolved by SDS–PAGE on Mini-PROTEAN TGX gels (Bio-Rad) at 150 V for 45 min and transferred to PVDF membranes using a Trans-Blot Turbo system (Bio-Rad). Membranes were blocked in 5% non-fat milk in TBST (Thermo Fisher Scientific) for 1 h at room temperature, then incubated overnight at 4 °C with primary antibodies diluted in 3% non-fat milk in TBST. To validate over-expression of LDHA, a mouse anti-FLAG (1:1000; Millipore Sigma F3165) primary antibody was used, and to validate over-expression of LDHC, a rabbit anti-FLAG primary antibody (1:400; Millipore Sigma F7425) was used. Following TBST washes, membranes were incubated with anti-rabbit HRP (Cytiva NA934) or anti-mouse HRP (Cytiva NA931) antibodies diluted 1:5000 in 5% non-fat milk in TBST for 1 hr at room temperature. Protein bands were detected using enhanced chemiluminescence (ECL; Thermo Fisher Scientific).

#### Enzyme activity assays

Recombinant human lactate dehydrogenase A (LDHA; Sigma-Aldrich SAE0049) was used for all enzymatic assays. The assay buffer consisted of 200 mM Tris-HCl supplemented with 0.2 mM dithiothreitol (DTT), adjusted to pH 8.0. NADH was prepared at a final concentration of 10 mM by dissolving 14.2 mg NADH in 2 mL of assay buffer. α-ketoisovalerate (αKIV) was prepared as a 600 mM stock solution and serially diluted two-fold to generate a 12-point concentration series. Reactions were assembled by combining 35 μL assay buffer, 5 μL LDHA (1 kU/mL), and 5 μL NADH in each well. Enzymatic reactions were initiated by the addition of 10 μL αKIV at the indicated concentrations. The rate of NADH consumption was quantified for 10 minutes on a plate reader using the absorbance of NADH at 340 nm.

#### Genotyping of *Bckdk* KO mice

Genomic DNA was isolated from mouse ear punch biopsies. Tissue was incubated in 25 mM NaOH and 0.2 mM EDTA at 100 °C for 1 hr and neutralized with 40 mM Tris-HCl (pH 7.5). Genotyping PCR was performed using allele-specific primers to detect wild-type and mutant Bdk alleles. The wild-type allele was amplified using forward primer AAGTGGAATAACTCACAGTAACTCAC and reverse primer CCTCGAGGATGCAGACGTGCTCAGC, yielding a 700 bp product. The mutant allele was amplified using forward primer AAATGGCGTTACTTAAGCTAGCTTGC and the same reverse primer, yielding a 400 bp product. PCR reactions (15 μL) were assembled using KOD Xtreme™ Hot Start DNA Polymerase (Sigma Aldrich) according to the manufacturer’s instructions, with 2 μL of genomic DNA per reaction. Amplification conditions were 94 °C for 2 min, followed by 35 cycles of 98 °C for 10 s, 55 °C for 45 s, and 68 °C for 15 s, with a final extension at 68 °C for 2 min. PCR products were resolved on 3% agarose gels in TAE buffer and visualized using SYBR Safe DNA stain.

#### Glucose tolerance tests

6- to 17-week-old *Bckdk*^+/+^, *Bckdk*^+/-^, and *Bckdk*^-/-^ mice were fasted beginning at 10:00 AM, and glucose tolerance tests (GTTs) were initiated at 1:00 PM. A sterile glucose solution was prepared at a concentration of 20 g/100 mL in water and administered intraperitoneally at a dose of 2 g/kg body weight, corresponding to an injection volume of 10 μL/g. Prior to glucose administration, mice were weighed and baseline blood glucose levels were measured. Blood glucose concentrations were subsequently assessed at 15, 30, 60, 90, and 120 minutes post-injection using a OneTouch Ultra Plus glucometer.

#### Tissue NADH/NAD^+^ and α-hydroxybutyrate measurements

Metabolites were extracted from flash-frozen tissue samples as previously described.^86^ Solvent A consisted of acetonitrile/methanol/water (40:40:20, v/v/v) supplemented with 0.1 M formic acid, and solvent B consisted of 15% (w/v) ammonium bicarbonate in water, maintained on ice.

Tissue samples were ground using a cryo-cup grinder. For every 20 mg of tissue, 1,000 µL of solvent A was added, followed by vortexing for 10 seconds and incubation on ice for 3 minutes. Solvent B (87 µL per 1,000 µL Solvent A) was then added, and samples were vortexed and incubated on dry ice for 20 minutes. Samples were centrifuged at 16,000 × g for 30 min at 4 °C, and 500 µL of the supernatant was collected and lyophilized overnight. Dried extracts were resuspended in 50 µL of acetonitrile/methanol/water (40:40:20, v/v/v) containing 10 mM ammonium bicarbonate. The resuspended samples were vortexed, mixed on a thermomixer at 4 °C, and incubated on ice for 20 minutes. Samples were then centrifuged at 16,000 × g for 20 min, and 40 µL of the clarified extract was transferred to autosampler vials for LC–MS analysis.

Samples were analyzed on an Orbitrap Exploris 240 high-resolution mass spectrometer (Thermo Fisher Scientific) equipped with an electrospray ionization source and coupled to a Vanquish Horizon UHPLC system. Polar metabolites (2 μL per sample) were separated using hydrophilic interaction chromatography on an iHILIC-(P) Classic column (2.1 × 150 mm, 5 µm; HILICON AB). The autosampler was maintained at 4 °C and the column temperature at 40 °C. Mobile phase A consisted of 20 mM ammonium bicarbonate in water, adjusted to pH 9.6 with ammonium hydroxide, while mobile phase B was acetonitrile. Separation was performed at a flow rate of 200 μL/min using the following gradient: 85 to 20% B from 0-18 minutes, hold at 20% B from 18-20 minutes, a linear increase from 20 to 85% B from 20-20.5 minutes, and hold at 85% B from 20.5-28 minutes. Each run was preceded by a 10-min equilibration at initial conditions.

The mass spectrometer was operated in full scan mode with polarity switching or in tSIM mode with negative polarity. In each mode, sheath, auxiliary, and sweep gas flows were set to 35, 10, and 0.5 units, respectively. The ion transfer tube temperature was maintained at 300 °C, and the vaporizer temperature was set to 35 °C. Spray voltages of 3.5 kV in positive mode and 3.25 kV in negative mode were used. In full scan mode, mass spectra were acquired over an m/z range of 70–1000 at a resolution of 60,000. Instrument parameters included an RF lens setting of 60%, an AGC target of 1 × 10⁷, and a maximum injection time of 100 ms. In tSIM mode, mass spectra were acquired at a resolution of 60,000, an RF lens of 60%, an AGC target of 1 × 10^6^, and a maximum injection time of 30 ms. In tSIM mode, the following compounds were detected (with retention time range in minutes): NAD^+^ (6.66-8.66) and NADH (4.59-8.59).

#### Blood chemistries

Approximately 90 μL of blood was obtained via tail blood sampling from 6- to 17-week-old *Bckdk*^+/+^, *Bckdk*^+/-^, and *Bckdk*^-/-^ mice. Samples were collected into K2EDTA-coated tubes (BD) and analyzed using an iSTAT handheld analyzer (Zoetis) with CHEM8+ cartridges (Zoetis).

#### Binge drinking model

Mice were fasted beginning at 7:30 AM. A 44.3% (v/v) solution of ethanol was prepared and delivered to mice via oral gavage at a dose of 3.5 g/kg. Control mice received an equivalent volume of water. Ethanol gavage was performed at 10:30 AM and repeated at 11:00 AM.

A branched-chain ketoacid (BCKA) mixture composed of 50 mg/mL of each of the following was prepared: 3-Methyl-2-oxobutanoic acid sodium salt (Thermo Fisher 189720010), (±)-3-Methyl-2-oxovaleric acid sodium salt (Sigma Aldrich K7125), α-Ketoisocaproic Acid (sodium salt) (Cayman Chemical 21052). A 10 μL/g dose of the BCKA mixture was administered intraperitoneally at 10:45 AM and again at 11:15 AM. Vehicle control mice received an equivalent volume of NaCl matched for total molarity (1.02 M).

At 4:30 PM, sub-mandibular blood was collected, after which mice were euthanized and livers were harvested for downstream analyses.

#### Multiple sequence alignment

Multiple sequence alignment of lactate dehydrogenase protein sequences were performed using NCBI COBALT.^87^ The resulting alignment was visualized using Jalview.^88^

#### Phylogenetic tree construction

The 100 closest sequences to the *Mus musculus* LDHC protein were curated with NCBI protein BLAST using the reference proteins database. The resulting sequences were assembled into a phylogenetic tree using the Jukes-Cantor genetic distance model and UPGMA tree build method in Geneious Prime 2026.0.1 (https://www.geneious.com). Species divergence was determined using TimeTree 5.^89^

#### Sperm NADH/NAD^+^ measurements

Mouse sperm were isolated, and capacitation was induced. Sperm from each mouse were split into different nutrient conditions during capacitation: glucose (5.5 mM) alone, glucose with sodium pyruvate (5 mM, Fisher Scientific AC132150250), glucose with BCKA sodium salts (1.67 mM each), glucose with sodium cyanide (10 mM, Millipore Sigma 380970) or rotenone (500 nM, Sigma Aldrich R8875), glucose with sodium cyanide or rotenone with sodium pyruvate, and glucose with sodium cyanide or rotenone with BCKAs.

Intracellular NADH/NAD⁺ ratios were quantified using the NAD/NADH-Glo Assay (Promega G9071) following manufacturer instructions. Capacitated sperm were transferred to microcentrifuge tubes and centrifuged at 300 × g for 5 min. The supernatant was aspirated, and cell pellets were resuspended in 60 µL phosphate-buffered saline (PBS). An equal volume (60 µL) of 0.2 N NaOH with 1% DTAB was added to initiate cell lysis. Samples were mixed on a thermomixer to ensure complete lysis and homogeneity. For differential quantification of NAD⁺ and NADH, 50 µL of each lysate was transferred to two separate wells of a 96-well plate. One set of wells was subjected to acid treatment for NADH degradation by addition of 25 µL of 0.4 N HCl per well, while the corresponding wells were processed in parallel. Plates were incubated at 60 °C for 15 min, followed by equilibration at room temperature for 10 min. Acid-treated samples were neutralized by addition of 25 µL of 0.5 M Trizma base per well, and 50 µL of 0.4 N HCl/0.5 M Trizma mixture was added to the other wells to equalize volume and buffer conditions. The NAD⁺/NADH-Glo detection reagent was prepared by combining 10 mL reconstituted luciferin detection reagent with 50 µL reductase, 50 µL reductase substrate, 50 µL NAD^+^ cycling enzyme, and 250 µL NAD^+^ cycling substrate. Detection reagent (100 µL) was added to each well, plates were shaken to mix, and luminescence was measured on a plate reader after incubation for 30–60 min.

#### [4-^2^H]glucose stable isotope tracer experiments

Mouse sperm were isolated and split into capacitation solution containing 5 mM [4-^2^H]glucose (Cambridge Isotope Laboratories DLM-9294) with 4 conditions: [4-^2^H]glucose only, [4-^2^H]glucose with 1 mM sodium pyruvate and 1 mM of each BCKA sodium salt, [4-^2^H]glucose with 10 mM sodium cyanide, and [4-^2^H]glucose with 10 mM sodium cyanide and 1 mM sodium pyruvate and 1 mM of each BCKA sodium salt.

After 3 hours of capacitation, sperm were transferred to microcentrifuge tubes and centrifuged at 300 × g for 5 min. The supernatant was saved for measurement of secreted electron acceptors. Metabolites were extracted by mixing 20 μL of liquid samples or standards with 200 μL of 80% methanol with 1 mM of the internal standard D8-Valine. The mixtures were incubated at −80 °C for 3 hours and vortexed. Next, the mixtures were centrifuged for 10 minutes at 16,000 × g at 4 °C. Supernatants were transferred to glass vials for storage or analysis.

#### Respirometry with Seahorse XF96

Mouse sperm bioenergetics were assayed using a Seahorse extracellular flux analyzer based on an established protocol.^90^ The day prior to the assay, Seahorse XF sensor cartridges were hydrated by adding 200 μL of Seahorse XF Calibrant to each well and incubating overnight at 37 °C in a non-CO_2_ incubator. Modified Tyrode’s (mT) medium was prepared using Milli-Q water containing NaCl (4.29 g/L), KCl (99.9 mg/L), MgCl_2_·6H_2_O (49.8 mg/L), NaH_2_PO_4_·2H_2_O (28.1 mg/L), CaCl_2_ (99.9 mg/L), HEPES (2.383 g/L), fatty acid-free BSA (4 mg/mL), and glucose (901 mg/L). The pH was adjusted to 7.4 at 37 °C, and the medium was sterile-filtered prior to use.

For plate coating, concanavalin A was dissolved at 5 mg/mL in water, aliquoted, and stored at −20 °C. Immediately before use, the stock was diluted to 0.5 mg/mL, and 20 μL was added to each well. Plates were incubated overnight at 4 °C. Prior to cell plating, wells were rinsed with 100 μL Milli-Q water, aspirated, and allowed to air-dry. Coated plates were used within 3 hours.

ETC inhibitor drugs were prepared for final assay concentrations of 5 μM oligomycin, 0.6 μM FCCP, 1 μM rotenone, and 1 μM antimycin A. Injectates (25 μL) were loaded into the appropriate ports of the hydrated sensor cartridge according to the experimental design. The loaded cartridge was incubated at 37 °C in a non-CO_2_ incubator for approximately 20 minutes before probe calibration.

Mouse sperm were isolated from 4 cauda epididymides in 2.2 mL of mT medium in one well of a 6-well plate. Sperm were split into 3 conditions: glucose only, glucose with 5 mM pyruvate, and glucose with 1.67 mM of each BCKA sodium salt. To attach sperm, the plate was centrifuged for 3.5 min at 1200 × g at 37 °C with 0 acceleration and 0 brake. This centrifugation step was repeated with the plate rotated 180 degrees.

Each mix-measure cycle on the Seahorse analyzer involved 2 minutes of mixing followed by 3 minutes of measurement. The analyzer was operated for 5 cycles at baseline, 3 cycles following oligomycin injection, 2 cycles following each FCCP injection, and 3 cycles following the injection of rotenone and antimycin.

ATP production rates were calculated from resulting oxygen consumption rate (OCR) and extracellular acidification rate (ECAR) values as previously described.^91^ ATP-linked OCR (OCR_ATP_) was calculated by subtracting average OCR during oligomycin treatment from the average basal OCR during the 4th and 5th measurements. Proton leak-linked OCR (OCR_Leak_) was calculated by subtracting average OCR during rotenone and antimycin treatment from the average OCR during oligomycin treatment. Total mitochondrial OCR (OCR_Mito_) was calculated by subtracting average OCR during rotenone and antimycin treatment from the average basal OCR during the 4th and 5th measurements. Basal proton production rate (H^+^_Basal_) was calculated by dividing the average basal ECAR during the 4th and 5th measurements by 0.175. Maximal proton production rate (H^+^_Max_) was calculated by average ECAR during oligomycin treatment by 0.175. The rate of ATP production from respiration (ATP_Respiration_) was calculated using the following equation:

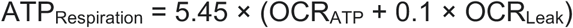

The rate of ATP production from fermentation during basal conditions and oligomycin-treated conditions was calculated according to the following equations:

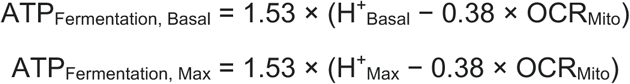

#### Respirometry with Oroboros

Respiration was measured using an Oroboros O_2_k respirometer (Oroboros Instruments). The 0.5-mL respiration chamber was equipped with a Clark-type oxygen electrode and maintained at 37 °C under continuous stirring. Calibration was performed at two oxygen concentrations: the saturated oxygen concentration under ambient air pressure (assumed to be 205 μM at 37 °C) and the zero-oxygen concentration following addition of dithionite (Na_2_S_2_O_4_). Oxygen concentration was recorded every 0.2 s. The oxygen consumption rate (OCR) was continuously determined and calculated as the slope of a linear fit over all measurements within the 8-s interval immediately preceding each OCR calculation. Non-mitochondrial respiration was corrected by subtracting the OCR measured after addition of mitochondrial inhibitors (1 μM rotenone and 2.5 μM antimycin A, abbreviated RA) at the end of each experiment. OCR signals were normalized to the basal OCR in each assay.

Sperm were isolated from the cauda epididymis of adult male mice by swim-out into human tubal fluid (HTF) medium lacking bicarbonate and bovine serum albumin (BSA), prepared with the following composition (mM): 102 NaCl, 4.7 KCl, 2.0 CaCl_2_, 0.37 KH2PO4, 0.2 MgSO4, and 20 HEPES. The medium was adjusted to pH 7.4 with NaOH and had an osmolarity of 320 ± 5 mOsm/L. Following swim-out, sperm concentration was adjusted to 10 × 10^6^ cells/mL, and 550 μL of the suspension was loaded into each chamber of the respirometer. After sperm loading, OCR was monitored for 5-10 min until a stable plateau was reached, at which point basal OCR was determined. Subsequently, either sodium salts of BCKAs (1.67 mM each of α-ketoisocaproic acid, (±)-3-methyl-2-oxovaleric acid, and α-ketoisovaleric acid), 5 mM sodium pyruvate, or 5 mM NaCl (as an osmotic control) was added. Oligomycin was then added to a final concentration of 1 μM, followed by stepwise titration of FCCP from 12.5 nM to 200 nM to assess maximal uncoupled respiration.

#### Mouse sperm motility quantification

Mouse sperm were isolated, and capacitation was induced. Sperm from each mouse were split into different nutrient conditions during capacitation: glucose (5.5 mM) alone, glucose with sodium pyruvate (5 mM), glucose with BCKA sodium salts (1.67 mM each), glucose with sodium cyanide (10 mM), glucose with sodium cyanide with sodium pyruvate, and glucose with sodium cyanide with BCKA sodium salts. 3 μL of capacitated mouse sperm from each experimental condition were loaded into pre-warmed (37 °C) 20-μm Leja four-chamber counting slides (IMV Technologies 025107-025108). Motility parameters were assessed using computer-assisted sperm analysis (CASA) with a Hamilton–Thorne digital image analyzer (HTR-CEROS II v1.7; Hamilton-Thorne Research).

Hyperactivated motility in mouse sperm was defined by a curvilinear velocity (VCL) exceeding 150 μm/s, an amplitude of lateral head displacement (ALH) greater than 7.0 μm, and a linearity (LIN) below 32% at a sampling rate of 60 Hz. The percentage of motile trajectories exhibiting hyperactive motility was calculated.

The following settings were used for imaging analysis:^92^ objective 1: Zeiss 10XNH; min total count: 200; frames acquired, 30; frame rate, 60 Hz; camera exposure: 8 ms; camera gain: 300; integrated time: 500 ms; elongation max%: 100; elongation min%: 1; head brightness min 170; head size max: 50 μm^2^; head size min: 5 μm^2^; static tail filter: false; tail brightness min: 70; tail brightness auto offset: 8; tail brightness mode: manual; progressive STR (%): 80; progressive VAP (μm/s): 25.

### Quantification and statistical analysis

Bar graphs depict mean ± SEM. Statistics were calculated using Prism 10 (GraphPad). Specific statistical tests are described in the figure legends.

## Notes

https://jain-lab-ucsf.github.io/hypoxia-metabolomics/

### Competing Interest Statement

The authors have declared no competing interest.

## REFERENCES

1. Mitchell, P. (1961). Coupling of Phosphorylation to Electron and Hydrogen Transfer by a Chemi-Osmotic type of Mechanism. Nature 191, 144–148. 10.1038/191144a0.

2. Sullivan, L.B., Gui, D.Y., Hosios, A.M., Bush, L.N., Freinkman, E., and Vander Heiden, M.G. (2015). Supporting Aspartate Biosynthesis Is an Essential Function of Respiration in Proliferating Cells. Cell 162, 552–563. 10.1016/j.cell.2015.07.017.

3. Birsoy, K., Wang, T., Chen, W.W., Freinkman, E., Abu-Remaileh, M., and Sabatini, D.M. (2015). An Essential Role of the Mitochondrial Electron Transport Chain in Cell Proliferation Is to Enable Aspartate Synthesis. Cell 162, 540–551. 10.1016/j.cell.2015.07.016.

4. Loscalzo, J. (2016). Adaptions to Hypoxia and Redox Stress. Circ. Res. 119, 511–513. 10.1161/CIRCRESAHA.116.309394.

5. Ge, M., Papagiannakopoulos, T., and Bar-Peled, L. (2024). Reductive stress in cancer: coming out of the shadows. Trends Cancer 10, 103–112. 10.1016/j.trecan.2023.10.002.

6. Xiao, W., and Loscalzo, J. (2020). Metabolic Responses to Reductive Stress. Antioxid. Redox Signal. 32, 1330–1347. 10.1089/ars.2019.7803.

7. Ido, Y., Kilo, C., and Williamson, J.R. (1997). Cytosolic NADH/NAD+, free radicals, and vascular dysfunction in early diabetes mellitus. Diabetologia 40, S115–S117. 10.1007/s001250051422.

8. Kim, W., Deik, A., Gonzalez, C., Gonzalez, M.E., Fu, F., Ferrari, M., Churchhouse, C.L., Florez, J.C., Jacobs, S.B.R., Clish, C.B., et al. (2019). Polyunsaturated Fatty Acid Desaturation Is a Mechanism for Glycolytic NAD+ Recycling. Cell Metab. 29, 856–870.e7. 10.1016/j.cmet.2018.12.023.

9. Jokinen, M.J., and Luukkonen, P.K. (2024). Hepatic mitochondrial reductive stress in the pathogenesis and treatment of steatotic liver disease. Trends Pharmacol. Sci. 45, 319–334. 10.1016/j.tips.2024.02.003.

10. Singh, C., Jin, B., Shrestha, N., Markhard, A.L., Panda, A., Calvo, S.E., Deik, A., Pan, X., Zuckerman, A.L., Saad, A.B., et al. (2024). ChREBP is activated by reductive stress and mediates GCKR-associated metabolic traits. Cell Metab. 36, 144–158.e7. 10.1016/j.cmet.2023.11.010.

11. Goodman, R.P., Markhard, A.L., Shah, H., Sharma, R., Skinner, O.S., Clish, C.B., Deik, A., Patgiri, A., Hsu, Y.-H.H., Masia, R., et al. (2020). Hepatic NADH reductive stress underlies common variation in metabolic traits. Nature 583, 122–126. 10.1038/s41586-020-2337-2.

12. Liu, S., Fu, S., Wang, G., Cao, Y., Li, L., Li, X., Yang, J., Li, N., Shan, Y., Cao, Y., et al. (2021). Glycerol-3-phosphate biosynthesis regenerates cytosolic NAD+ to alleviate mitochondrial disease. Cell Metab. 33, 1974–1987.e9. 10.1016/j.cmet.2021.06.013.

13. Spinelli, J.B., Rosen, P.C., Sprenger, H.-G., Puszynska, A.M., Mann, J.L., Roessler, J.M., Cangelosi, A.L., Henne, A., Condon, K.J., Zhang, T., et al. (2021). Fumarate is a terminal electron acceptor in the mammalian electron transport chain. Science 374, 1227–1237. 10.1126/science.abi7495.

14. Chouchani, E.T., Pell, V.R., Gaude, E., Aksentijević, D., Sundier, S.Y., Robb, E.L., Logan, A., Nadtochiy, S.M., Ord, E.N.J., Smith, A.C., et al. (2014). Ischaemic accumulation of succinate controls reperfusion injury through mitochondrial ROS. Nature 515, 431–435. 10.1038/nature13909.

15. Hochachka, P.W., and Somero, G.N. (2002). Biochemical Adaptation: Mechanism and Process in Physiological Evolution (Oxford University Press) 10.1093/oso/9780195117028.001.0001.

16. Firth, J.D., Ebert, B.L., and Ratcliffe, P.J. (1995). Hypoxic regulation of lactate dehydrogenase A. Interaction between hypoxia-inducible factor 1 and cAMP response elements. J. Biol. Chem. 270, 21021–21027. 10.1074/jbc.270.36.21021.

17. Papandreou, I., Cairns, R.A., Fontana, L., Lim, A.L., and Denko, N.C. (2006). HIF-1 mediates adaptation to hypoxia by actively downregulating mitochondrial oxygen consumption. Cell Metab. 3, 187–197. 10.1016/j.cmet.2006.01.012.

18. Epstein, A.C., Gleadle, J.M., McNeill, L.A., Hewitson, K.S., O’Rourke, J., Mole, D.R., Mukherji, M., Metzen, E., Wilson, M.I., Dhanda, A., et al. (2001). C. elegans EGL-9 and mammalian homologs define a family of dioxygenases that regulate HIF by prolyl hydroxylation. Cell 107, 43–54. 10.1016/s0092-8674(01)00507-4.

19. Ivan, M., Kondo, K., Yang, H., Kim, W., Valiando, J., Ohh, M., Salic, A., Asara, J.M., Lane, W.S., and Kaelin Jr., W.G. (2001). HIFα Targeted for VHL-Mediated Destruction by Proline Hydroxylation: Implications for O 2 Sensing. Science 292, 464–468. 10.1126/science.1059817.

20. Jaakkola, P., Mole, D.R., Tian, Y.M., Wilson, M.I., Gielbert, J., Gaskell, S.J., von Kriegsheim, A., Hebestreit, H.F., Mukherji, M., Schofield, C.J., et al. (2001). Targeting of HIF-alpha to the von Hippel-Lindau ubiquitylation complex by O_2_-regulated prolyl hydroxylation. Science 292, 468–472. 10.1126/science.1059796.

21. Shoubridge, E.A., and Hochachka, P.W. (1980). Ethanol: Novel End Product of Vertebrate Anaerobic Metabolism. Science 209, 308–309. 10.1126/science.7384807.

22. Van den Thillart, G., Van Berge-Henegouwen, M., and Kesbeke, F. (1983). Anaerobic metabolism of goldfish, *Carassius auratus* (L.): Ethanol and CO_2_ excretion rates and anoxia tolerance at 20, 10 and 5°C. Comp. Biochem. Physiol. A Physiol. 76, 295–300. 10.1016/0300-9629(83)90330-4.

23. van den Thillart, G., van Waarde, A., Dobbe, F., and Kesbeke, F. (1982). Anaerobic energy metabolism of goldfish,Carassius auratus (L.). J. Comp. Physiol. 146, 41–49. 10.1007/bf00688715.

24. Collicutt, J.M., and Hochachka, P.W. (1977). The anaerobic oyster heart: Coupling of glucose and aspartate fermentation. J. Comp. Physiol. 115, 147–157. 10.1007/BF00692526.

25. Grieshaber, M.K., Hardewig, I., Kreutzer, U., and Pörtner, H.-O. (1994). Physiological and metabolic responses to hypoxia in invertebrates. In Reviews of Physiology, Biochemistry and Pharmacology, Volume 125: Volume: 125 (Springer), pp. 43–147. 10.1007/BFb0030909.

26. Westbrook, R.L., Bridges, E., Roberts, J., Escribano-Gonzalez, C., Eales, K.L., Vettore, L.A., Walker, P.D., Vera-Siguenza, E., Rana, H., Cuozzo, F., et al. (2022). Proline synthesis through PYCR1 is required to support cancer cell proliferation and survival in oxygen-limiting conditions. Cell Rep. 38, 110320. 10.1016/j.celrep.2022.110320.

27. Intlekofer, A.M., Dematteo, R.G., Venneti, S., Finley, L.W.S., Lu, C., Judkins, A.R., Rustenburg, A.S., Grinaway, P.B., Chodera, J.D., Cross, J.R., et al. (2015). Hypoxia Induces Production of L-2-Hydroxyglutarate. Cell Metab. 22, 304–311. 10.1016/j.cmet.2015.06.023.

28. Oldham, W.M., Clish, C., Yang, Y., and Loscalzo, J. (2015). Hypoxia-mediated Increases in L-2-hydroxyglutarate Coordinate the Metabolic Response to Reductive Stress. Cell Metab. 22, 291–303. 10.1016/j.cmet.2015.06.021.

29. Intlekofer, A.M., Wang, B., Liu, H., Shah, H., Carmona-Fontaine, C., Rustenburg, A.S., Salah, S., Gunner, M.R., Chodera, J.D., Cross, J.R., et al. (2017). L-2-hydroxyglutarate production arises from non-canonical enzyme function at acidic pH. Nat. Chem. Biol. 13, 494–500. 10.1038/nchembio.2307.

30. Hui, S., Ghergurovich, J.M., Morscher, R.J., Jang, C., Teng, X., Lu, W., Esparza, L.A., Reya, T., Le Zhan, null, Yanxiang Guo, J., et al. (2017). Glucose feeds the TCA cycle via circulating lactate. Nature 551, 115–118. 10.1038/nature24057.

31. Rabinowitz, J.D., and Enerbäck, S. (2020). Lactate: the ugly duckling of energy metabolism. Nat. Metab. 2, 566–571. 10.1038/s42255-020-0243-4.

32. Okuda, S., Yamada, T., Hamajima, M., Itoh, M., Katayama, T., Bork, P., Goto, S., and Kanehisa, M. (2008). KEGG Atlas mapping for global analysis of metabolic pathways. Nucleic Acids Res. 36, W423–W426. 10.1093/nar/gkn282.

33. Sullivan, L.B., Luengo, A., Danai, L.V., Bush, L.N., Diehl, F.F., Hosios, A.M., Lau, A.N., Elmiligy, S., Malstrom, S., Lewis, C.A., et al. (2018). Aspartate is an endogenous metabolic limitation for tumour growth. Nat. Cell Biol. 20, 782–788. 10.1038/s41556-018-0125-0.

34. Hummel, W., Schütte, H., and Kula, M.-R. (1985). d-2-hydroxyisocaproate dehydrogenase from Lactobacillus casei. Appl. Microbiol. Biotechnol. 21, 7–15. 10.1007/BF00252354.

35. Schütte, H., Hummel, W., and Kula, M.-R. (1984). l-2-hydroxyisocaproate dehydrogenase— A new enzyme from Lactobacillus confusus for the stereospecific reduction of 2-ketocarboxylic acids. Appl. Microbiol. Biotechnol. 19, 167–176. 10.1007/BF00256449.

36. Kim, J., Darley, D., Selmer, T., and Buckel, W. (2006). Characterization of (R)-2-Hydroxyisocaproate Dehydrogenase and a Family III Coenzyme A Transferase Involved in Reduction of l-Leucine to Isocaproate by Clostridium difficile. Appl. Environ. Microbiol. 72, 6062–6069. 10.1128/AEM.00772-06.

37. Neinast, M., Murashige, D., and Arany, Z. (2019). Branched Chain Amino Acids. Annu. Rev. Physiol. 81, 139–164. 10.1146/annurev-physiol-020518-114455.

38. Villani, G.R., Gallo, G., Scolamiero, E., Salvatore, F., and Ruoppolo, M. (2017). “Classical organic acidurias”: diagnosis and pathogenesis. Clin. Exp. Med. 17, 305–323. 10.1007/s10238-016-0435-0.

39. Daniel, N., Nachbar, R.T., Tran, T.T.T., Ouellette, A., Varin, T.V., Cotillard, A., Quinquis, L., Gagné, A., St-Pierre, P., Trottier, J., et al. (2022). Gut microbiota and fermentation-derived branched chain hydroxy acids mediate health benefits of yogurt consumption in obese mice. Nat. Commun. 13, 1343. 10.1038/s41467-022-29005-0.

40. Wallace, M., Green, C.R., Roberts, L.S., Lee, Y.M., McCarville, J.L., Sanchez-Gurmaches, J., Meurs, N., Gengatharan, J.M., Hover, J.D., Phillips, S.A., et al. (2018). Enzyme promiscuity drives branched-chain fatty acid synthesis in adipose tissues. Nat. Chem. Biol. 14, 1021–1031. 10.1038/s41589-018-0132-2.

41. Green, C.R., Wallace, M., Divakaruni, A.S., Phillips, S.A., Murphy, A.N., Ciaraldi, T.P., and Metallo, C.M. (2016). Branched-chain amino acid catabolism fuels adipocyte differentiation and lipogenesis. Nat. Chem. Biol. 12, 15–21. 10.1038/nchembio.1961.

42. Yin, X., Chan, L.S., Bose, D., Jackson, A.U., VandeHaar, P., Locke, A.E., Fuchsberger, C., Stringham, H.M., Welch, R., Yu, K., et al. (2022). Genome-wide association studies of metabolites in Finnish men identify disease-relevant loci. Nat. Commun. 13, 1644. 10.1038/s41467-022-29143-5.

43. Heemskerk, M.M., van Harmelen, V.J., van Dijk, K.W., and van Klinken, J.B. (2016). Reanalysis of mGWAS results and in vitro validation show that lactate dehydrogenase interacts with branched-chain amino acid metabolism. Eur. J. Hum. Genet. 24, 142–145. 10.1038/ejhg.2015.106.

44. Kwong, A., Boughton, A.P., Wang, M., VandeHaar, P., Boehnke, M., Abecasis, G., and Kang, H.M. (2022). FIVEx: an interactive eQTL browser across public datasets. Bioinforma. Oxf. Engl. 38, 559–561. 10.1093/bioinformatics/btab614.

45. Titov, D.V., Cracan, V., Goodman, R.P., Peng, J., Grabarek, Z., and Mootha, V.K. (2016). Complementation of mitochondrial electron transport chain by manipulation of the NAD+/NADH ratio. Science 352, 231–235. 10.1126/science.aad4017.

46. Borst, P. (2020). The malate-aspartate shuttle (Borst cycle): How it started and developed into a major metabolic pathway. IUBMB Life 72, 2241–2259. 10.1002/iub.2367.

47. Hu, Q., Wu, D., Walker, M., Wang, P., Tian, R., and Wang, W. (2021). Genetically encoded biosensors for evaluating NAD+/NADH ratio in cytosolic and mitochondrial compartments. Cell Rep. Methods 1, 100116. 10.1016/j.crmeth.2021.100116.

48. Verkerke, A.R.P., Wang, D., Yoshida, N., Taxin, Z.H., Shi, X., Zheng, S., Li, Y., Auger, C., Oikawa, S., Yook, J.-S., et al. (2024). BCAA-nitrogen flux in brown fat controls metabolic health independent of thermogenesis. Cell 0. 10.1016/j.cell.2024.03.030.

49. Jang, C., Oh, S.F., Wada, S., Rowe, G.C., Liu, L., Chan, M.C., Rhee, J., Hoshino, A., Kim, B., Ibrahim, A., et al. (2016). A branched-chain amino acid metabolite drives vascular fatty acid transport and causes insulin resistance. Nat. Med. 22, 421–426. 10.1038/nm.4057.

50. Parker, P.J., and Randle, P.J. (1978). Partial purification and properties of branched-chain 2-oxo acid dehydrogenase of ox liver. Biochem. J. 171, 751–757. 10.1042/bj1710751.

51. Joshi, M.A., Jeoung, N.H., Obayashi, M., Hattab, E.M., Brocken, E.G., Liechty, E.A., Kubek, M.J., Vattem, K.M., Wek, R.C., and Harris, R.A. (2006). Impaired growth and neurological abnormalities in branched-chain α-keto acid dehydrogenase kinase-deficient mice. Biochem. J. 400, 153–162. 10.1042/BJ20060869.

52. Neinast, M.D., Jang, C., Hui, S., Murashige, D.S., Chu, Q., Morscher, R.J., Li, X., Zhan, L., White, E., Anthony, T.G., et al. (2019). Quantitative analysis of the whole-body metabolic fate of branched chain amino acids. Cell Metab. 29, 417–429.e4. 10.1016/j.cmet.2018.10.013.

53. Morville, T., Sahl, R.E., Moritz, T., Helge, J.W., and Clemmensen, C. (2020). Plasma Metabolome Profiling of Resistance Exercise and Endurance Exercise in Humans. Cell Rep. 33, 108554. 10.1016/j.celrep.2020.108554.

54. Páez-Franco, J.C., Torres-Ruiz, J., Sosa-Hernández, V.A., Cervantes-Díaz, R., Romero-Ramírez, S., Pérez-Fragoso, A., Meza-Sánchez, D.E., Germán-Acacio, J.M., Maravillas-Montero, J.L., Mejía-Domínguez, N.R., et al. (2021). Metabolomics analysis reveals a modified amino acid metabolism that correlates with altered oxygen homeostasis in COVID-19 patients. Sci. Rep. 11, 6350. 10.1038/s41598-021-85788-0.

55. Zheng, Y., Yu, B., Alexander, D., Steffen, L.M., Nettleton, J.A., and Boerwinkle, E. (2014). Metabolomic patterns and alcohol consumption in African Americans in the Atherosclerosis Risk in Communities Study123. Am. J. Clin. Nutr. 99, 1470–1478. 10.3945/ajcn.113.074070.

56. Goldberg, E., Eddy, E.M., Duan, C., and Odet, F. (2010). LDHC: The Ultimate Testis-Specific Gene. J. Androl. 31, 86–94. 10.2164/jandrol.109.008367.

57. Blanco, A., Burgos, C., Gerez de Burgos, N.M., and Montamat, E.E. (1976). Properties of the testicular lactate dehydrogenase isoenzyme. Biochem. J. 153, 165–172.

58. Madern, D. (2002). Molecular evolution within the L-malate and L-lactate dehydrogenase super-family. J. Mol. Evol. 54, 825–840. 10.1007/s00239-001-0088-8.

59. Chambellon, E., Rijnen, L., Lorquet, F., Gitton, C., van Hylckama Vlieg, J.E.T., Wouters, J.A., and Yvon, M. (2009). The d-2-Hydroxyacid Dehydrogenase Incorrectly Annotated PanE Is the Sole Reduction System for Branched-Chain 2-Keto Acids in Lactococcus lactis. J. Bacteriol. 191, 873–881. 10.1128/JB.01114-08.

60. Domenech, J., and Ferrer, J. (2006). A new d-2-hydroxyacid dehydrogenase with dual coenzyme-specificity from *Haloferax mediterranei*, sequence analysis and heterologous overexpression. Biochim. Biophys. Acta BBA - Gen. Subj. 1760, 1667–1674. 10.1016/j.bbagen.2006.08.024.

61. Coronel, C., Rovai, L.E., Gerez de Burgos, N.M., Burgos, C., and Blanco, A. (1981). Properties of α-hydroxyacid dehydrogenase isozymes from *Trypanosoma cruzi*. Mol. Biochem. Parasitol. 4, 29–38. 10.1016/0166-6851(81)90026-8.

62. Westrop, G.D., Wang, L., Blackburn, G.J., Zhang, T., Zheng, L., Watson, D.G., and Coombs, G.H. (2017). Metabolomic profiling and stable isotope labelling of Trichomonas vaginalis and Tritrichomonas foetus reveal major differences in amino acid metabolism including the production of 2-hydroxyisocaproic acid, cystathionine and S-methylcysteine. PLOS ONE 12, e0189072. 10.1371/journal.pone.0189072.

63. Elizondo, S., Chena, M.A., Rodríguez-Páez, L., Nogueda, B., Baeza, I., and Wong, C. (2003). Inhibition of Trypanosoma cruzi α-Hydroxyacid Dehydrogenase-isozyme II by N-Isopropyl Oxamate and its Effect on Intact Epimastigotes. J. Enzyme Inhib. Med. Chem. 18, 265–271. 10.1080/1475636031000071826.

64. Montamat, E.E., Burgos, C., Gerez de Burgos, N.M., Rovai, L.E., Blanco, A., and Segura, E.L. (1982). Inhibitory Action of Gossypol on Enzymes and Growth of Trypanosoma cruzi. Science 218, 288–289. 10.1126/science.6750791.

65. Odet, F., Duan, C., Willis, W.D., Goulding, E.H., Kung, A., Eddy, E.M., and Goldberg, E. (2008). Expression of the gene for mouse lactate dehydrogenase C (Ldhc) is required for male fertility. Biol. Reprod. 79, 26–34. 10.1095/biolreprod.108.068353.

66. Niu, X., Stancliffe, E., Gelman, S.J., Wang, L., Schwaiger-Haber, M., Rowles, J.L., Shriver, L.P., and Patti, G.J. (2023). Cytosolic and mitochondrial NADPH fluxes are independently regulated. Nat. Chem. Biol., 1–9. 10.1038/s41589-023-01283-9.

67. Lewis, C.A., Parker, S.J., Fiske, B.P., McCloskey, D., Gui, D.Y., Green, C.R., Vokes, N.I., Feist, A.M., Vander Heiden, M.G., and Metallo, C.M. (2014). Tracing Compartmentalized NADPH Metabolism in the Cytosol and Mitochondria of Mammalian Cells. Mol. Cell 55, 253–263. 10.1016/j.molcel.2014.05.008.

68. Suarez, S.S. (2008). Control of hyperactivation in sperm. Hum. Reprod. Update 14, 647– 657. 10.1093/humupd/dmn029.

69. Schmidt, C.A., Hale, B.J., Bhowmick, D., Miller, W.J., Neufer, P.D., and Geyer, C.B. (2024). Pyruvate modulation of redox potential controls mouse sperm motility. Dev. Cell 59, 79–90.e6. 10.1016/j.devcel.2023.11.011.

70. Miki, K., Qu, W., Goulding, E.H., Willis, W.D., Bunch, D.O., Strader, L.F., Perreault, S.D., Eddy, E.M., and O’Brien, D.A. (2004). Glyceraldehyde 3-phosphate dehydrogenase-S, a sperm-specific glycolytic enzyme, is required for sperm motility and male fertility. Proc. Natl. Acad. Sci. 101, 16501–16506. 10.1073/pnas.0407708101.

71. Fraser, L.R., and Quinn, P.J. (1981). A glycolytic product is obligatory for initiation of the sperm acrosome reaction and whiplash motility required for fertilization in the mouse. J. Reprod. Fertil. 61, 25–35. 10.1530/jrf.0.0610025.

72. Krisfalusi, M., Miki, K., Magyar, P.L., and O’Brien, D.A. (2006). Multiple glycolytic enzymes are tightly bound to the fibrous sheath of mouse spermatozoa. Biol. Reprod. 75, 270–278. 10.1095/biolreprod.105.049684.

73. Bunch, D. 0., Welch, J.E., Magyar, P.L., Eddy, E.M., and O’Brien, D.A. (1998). Glyceraldehyde 3-Phosphate Dehydrogenase-S Protein Distribution during Mouse Spermatogenesis1. Biol. Reprod. 58, 834–841. 10.1095/biolreprod58.3.834.

74. Cummins, J. (2009). Sperm motility and energetics. In Sperm Biology (Academic Press), pp. 185–206. 10.1016/B978-0-12-372568-4.00005-7.

75. Turner, R.M. (2005). Moving to the beat: a review of mammalian sperm motility regulation. Reprod. Fertil. Dev. 18, 25–38. 10.1071/RD05120.

76. Mukai, C., and Okuno, M. (2004). Glycolysis Plays a Major Role for Adenosine Triphosphate Supplementation in Mouse Sperm Flagellar Movement. Biol. Reprod. 71, 540–547. 10.1095/biolreprod.103.026054.

77. Stadtman, T.C., Elliott, P., and Tiemann, L. (1958). Studies on the enzymic reduction of amino acids. III. Phosphate esterification coupled with glycine reduction. J. Biol. Chem. 231, 961–973.

78. Stickland, L.H. (1934). Studies in the metabolism of the strict anaerobes (genus Clostridium). Biochem. J. 28, 1746–1759. 10.1042/bj0281746.

79. Clarke, P.H., and Elsden, S.R. (1980). The earliest catabolic pathways. J. Mol. Evol. 15, 333–338. 10.1007/BF01733139.

80. Livingstone, D.R. (1991). Origins and Evolution of Pathways of Anaerobic Metabolism in the Animal Kingdom. Am. Zool. 31, 522–534. 10.1093/icb/31.3.522.

81. Darszon, A., Nishigaki, T., López-González, I., Visconti, P.E., and Treviño, C.L. (2020). Differences and Similarities: The Richness of Comparative Sperm Physiology. Physiol. Bethesda Md 35, 196–208. 10.1152/physiol.00033.2019.

82. Jr, J.S.Z., Hodgkinson, C.A., Wright, M., Klise, A., Sundin, O., Broman, K.W., Hejtmancik, F., Huang, H., Patek, B., Sergeev, Y., et al. (2016). A Spontaneous Missense Mutation in Branched Chain Keto Acid Dehydrogenase Kinase in the Rat Affects Both the Central and Peripheral Nervous Systems. PLOS ONE 11, e0160447. 10.1371/journal.pone.0160447.

83. Shamailova, Y., Farooq, S.A., Gilmore, M.E., Stanek, T.J., Lopez, E.M., Wetstein, B.B., Mirek, E.T., Anthony, T.G., and Snyder, E.M. (2025). Branched chain amino acid sufficiency is necessary for proper luteinizing hormone response and testosterone synthesis. Reprod. Biol. 26, 101094. 10.1016/j.repbio.2025.101094.

84. Doench, J.G., Fusi, N., Sullender, M., Hegde, M., Vaimberg, E.W., Donovan, K.F., Smith, I., Tothova, Z., Wilen, C., Orchard, R., et al. (2016). Optimized sgRNA design to maximize activity and minimize off-target effects of CRISPR-Cas9. Nat. Biotechnol. 34, 184–191. 10.1038/nbt.3437.

85. Sanson, K.R., Hanna, R.E., Hegde, M., Donovan, K.F., Strand, C., Sullender, M.E., Vaimberg, E.W., Goodale, A., Root, D.E., Piccioni, F., et al. (2018). Optimized libraries for CRISPR-Cas9 genetic screens with multiple modalities. Nat. Commun. 9, 5416. 10.1038/s41467-018-07901-8.

86. Lu, W., Wang, L., Chen, L., Hui, S., and Rabinowitz, J.D. (2018). Extraction and Quantitation of Nicotinamide Adenine Dinucleotide Redox Cofactors. Antioxid. Redox Signal. 28, 167–179. 10.1089/ars.2017.7014.

87. Papadopoulos, J.S., and Agarwala, R. (2007). COBALT: constraint-based alignment tool for multiple protein sequences. Bioinformatics 23, 1073–1079. 10.1093/bioinformatics/btm076.

88. Waterhouse, A.M., Procter, J.B., Martin, D.M.A., Clamp, M., and Barton, G.J. (2009). Jalview Version 2—a multiple sequence alignment editor and analysis workbench. Bioinformatics 25, 1189–1191. 10.1093/bioinformatics/btp033.

89. Kumar, S., Suleski, M., Craig, J.M., Kasprowicz, A.E., Sanderford, M., Li, M., Stecher, G., and Hedges, S.B. (2022). TimeTree 5: An Expanded Resource for Species Divergence Times. Mol. Biol. Evol. 39, msac174. 10.1093/molbev/msac174.

90. Roldan, E.R.S., Tourmente, M., Sanchez-Rodriguez, A., and Rial, E. (2025). Bioenergetics of Rodent Spermatozoa. In Spermatology: Methods and Protocols, M. Álvarez-Rodríguez, ed. (Springer US), pp. 267–288. 10.1007/978-1-0716-4406-5_19.

91. Desousa, B.R., Kim, K.K., Jones, A.E., Ball, A.B., Hsieh, W.Y., Swain, P., Morrow, D.H., Brownstein, A.J., Ferrick, D.A., Shirihai, O.S., et al. (2023). Calculation of ATP production rates using the Seahorse XF Analyzer. EMBO Rep. 24, e56380. 10.15252/embr.202256380.

92. Molina, L.C.P., Gunderson, S., Riley, J., Lybaert, P., Borrego-Alvarez, A., Jungheim, E.S., and Santi, C.M. (2020). Membrane Potential Determined by Flow Cytometry Predicts Fertilizing Ability of Human Sperm. Front. Cell Dev. Biol. 7, 387. 10.3389/fcell.2019.00387.

