## Supplemental Figures for "Branched-chain amino acid fermentation as an alternative mammalian electron sink"

### SUPPLEMENTARY FIGURES

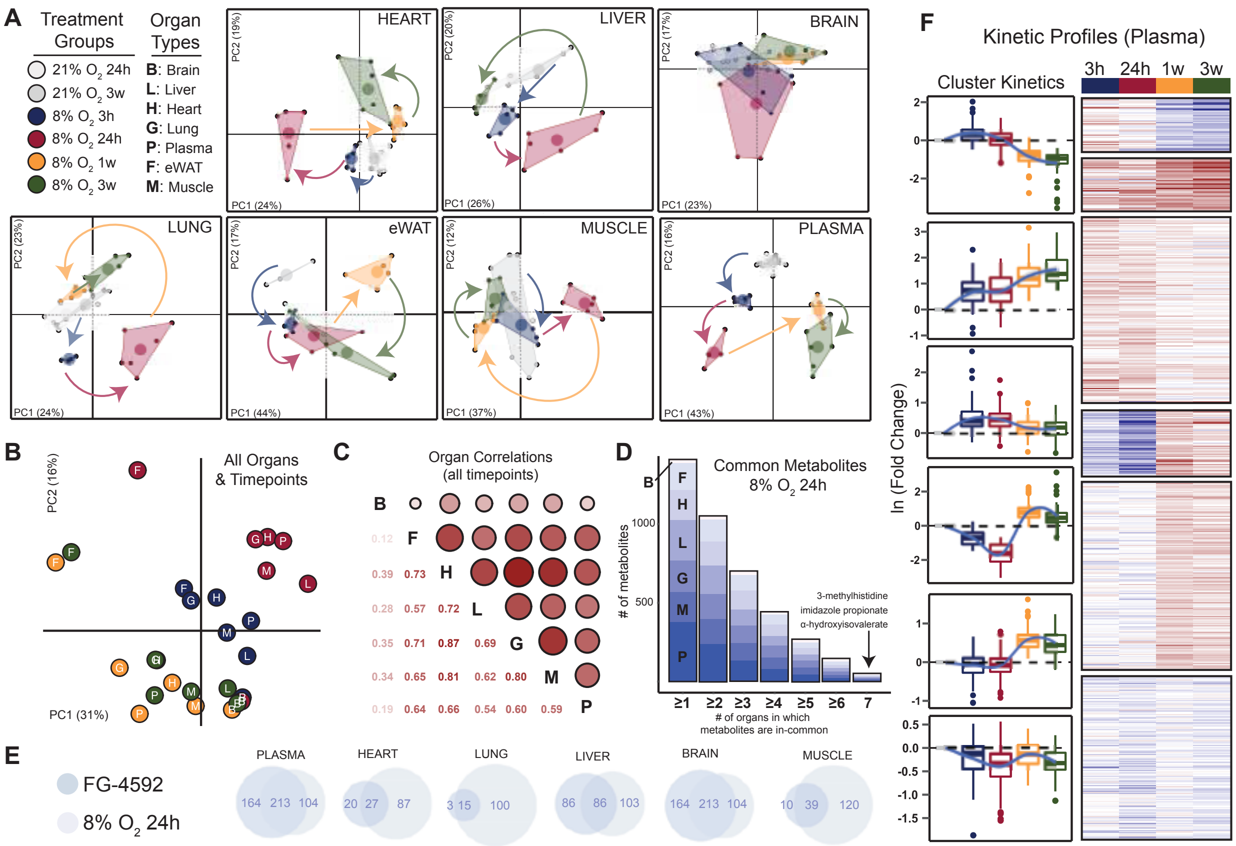


**Figure S1. Untargeted Metabolomics of Hypoxic Mouse Organs**
(A) Organ-specific principal component analysis (PCA) of metabolic profiles from mice housed at 21% $O_2$ and 8% $O_2$ for different amounts of time, using the most variable metabolites (variance > 0.5).
(B) Cross-organ PCA of hypoxia-responsive metabolites, classified as the compounds that were significantly altered over time (FDR ≤ 0.05) in at least 5 out of 7 organs. Analysis used linear model coefficients for each organ-timepoint combination under 8% $O_2$ exposure.
(C) Inter-organ correlation of hypoxia-responsive metabolites. Pair-wise correlations were calculated using linear model coefficients for hypoxia-responsive metabolites. Circle size and color intensity reflect correlation strength.
(D) Number of metabolites significantly different between 8% $O_2$ 24h and 21% $O_2$ 24h (p ≤ 0.05) in one or more organs. Bars show cumulative counts of metabolites altered in at least 1 organ up to the metabolites altered in all 7 organs.
(E) Venn diagrams comparing metabolites significantly altered (FDR ≤ 0.05) by pharmacological HIF stabilization (FG-4592) versus environmental hypoxia (8% $O_2$ 24h), each relative to 21% $O_2$ 24h controls.
(F) Clustered heatmap of plasma metabolites significantly altered over time under 8% $O_2$ exposure (FDR ≤ 0.05), grouped by k-means clustering (k = 6), alongside the distribution of cluster-specific effect sizes across timepoints.

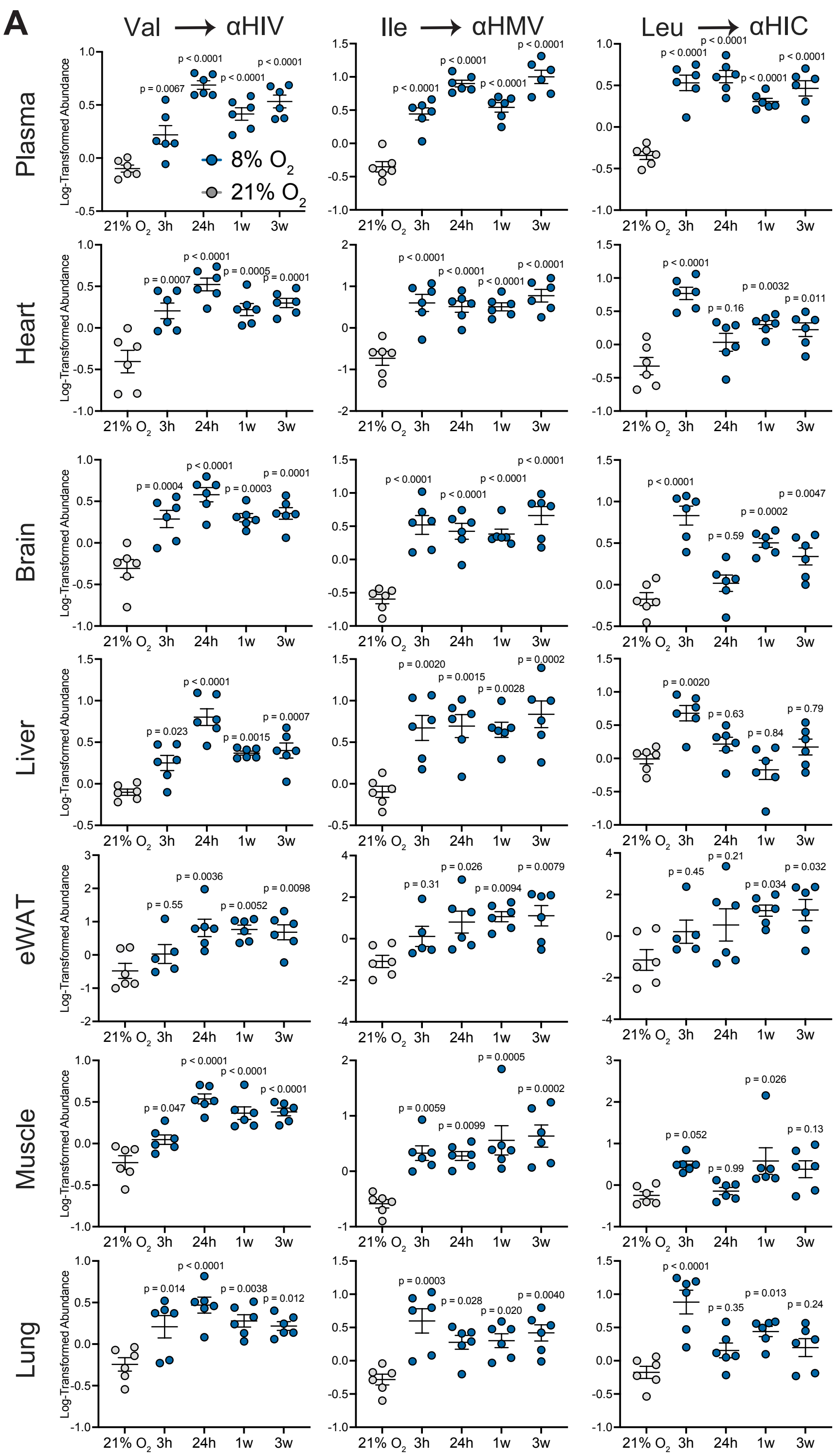


**Figure S2. Branched-chain hydroxyacid abundance by tissue**

(A) Log-transformed abundance of each BCHA in tissues from mice at 21% $O_2$ or from mice exposed to 8% $O_2$ for 3 hours, 24 hours, 1 week or 3 weeks.

Mean ± SEM are shown. Significance was determined by one-way ANOVA followed by Tukey's multiple comparisons test.

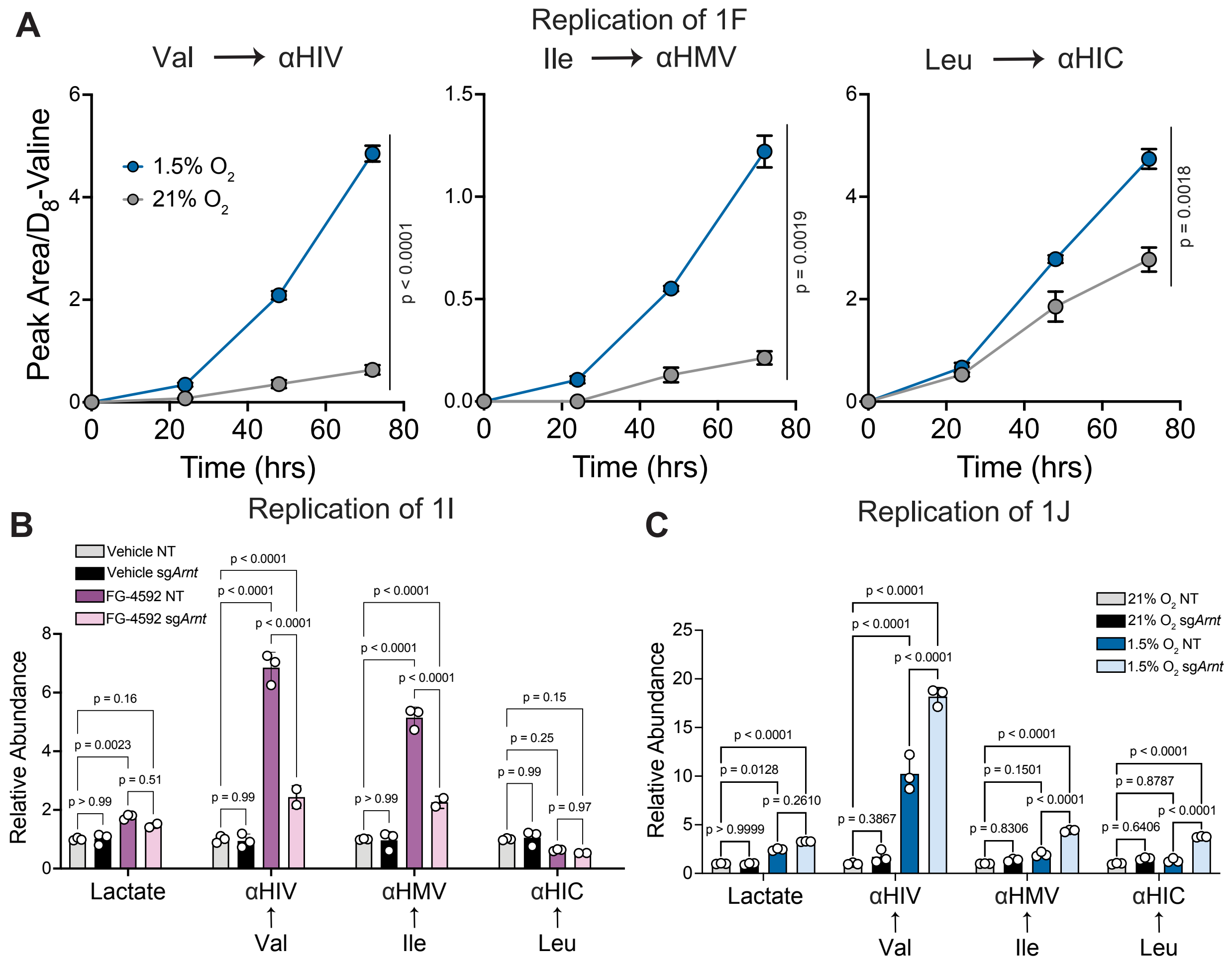


**Figure S3. Hypoxia and HIF activation promote BCAA fermentation**

(A) Abundance of $^{13}$C-BCHAs in conditioned media from 3T3-L1 adipocytes cultured at 21% $O_2$ or 1.5% $O_2$ for 24, 48, and 72 hours. n = 3 wells.

(B-C) Relative abundance of $^{13}$C-BCHAs in conditioned media from control or *Arnt*-deficient 3T3-L1 adipocytes after 3-day treatment with 0.15% DMSO or 75 µM FG-4592 (B) or 3-day treatment with 21% O2 or 1.5% O2 (C).

Mean ± SD are shown. Significance was determined by two-way ANOVA followed by Šídák's multiple comparisons test (A) or Tukey's multiple comparisons test (B-C).

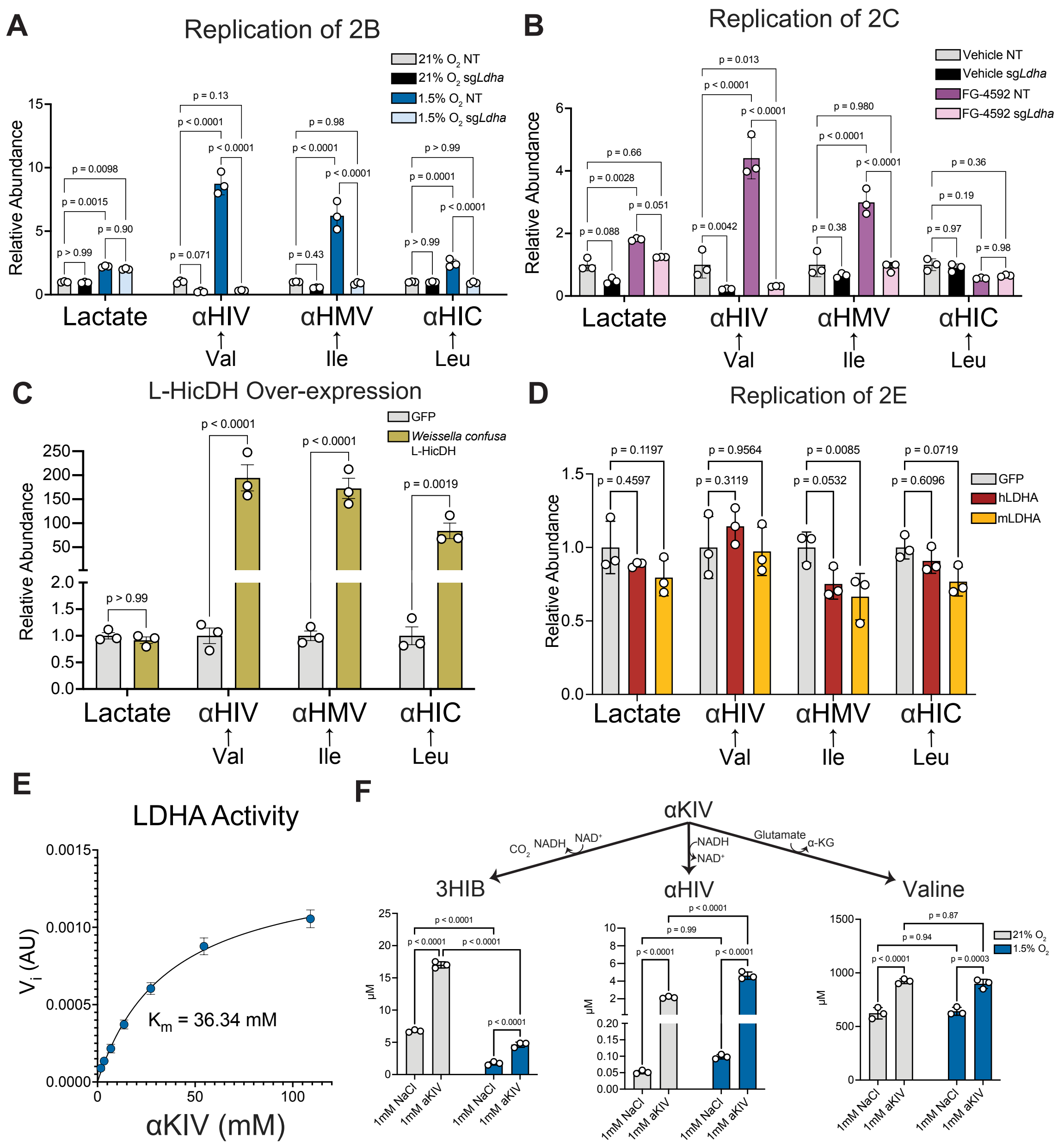


**Figure S4. Enzymatic regulation of BCAA fermentation *in vitro*.**

(A-B) Relative abundance of lactate and [13]C-BCHAs in conditioned media from control or *Ldha*-deficient 3T3-L1 adipocytes after 3-day treatment with 21% $O_2$ or 1.5% $O_2$ (A) or 3-day treatment with 0.15% DMSO or 75 µM FG-4592 (B). n = 3 wells.

(C) Relative abundance of lactate and [13]C-BCHAs in conditioned media from 3T3-L1 adipocytes overexpressing GFP or *Weissella confusa* L-hydroxyisocaproate dehydrogenase after 3 days of incubation at 21% $O_2$. n = 3 wells.

(D) Relative abundance of lactate and [13]C-BCHAs in conditioned media from 3T3-L1 adipocytes overexpressing GFP, mouse LDHA, or human LDHA after 3 days of incubation at 21% $O_2$. n = 3 wells.

(E) LDHA activity measured as the initial rate of change in NADH absorbance at different concentrations of α-ketoisovalerate (αKIV). n = 3 replicates.

(F) Concentration of valine metabolites in conditioned media from wild-type 3T3-L1 adipocytes supplemented with 1 mM NaCl or 1 mM αKIV and cultured in 21% $O_2$ or 1.5% $O_2$ for 1 day.

Mean ± SD are shown. Significance was determined by two-way ANOVA followed by Tukey's multiple comparisons test (A-B, D, F) or Šídák's multiple comparisons test (C).

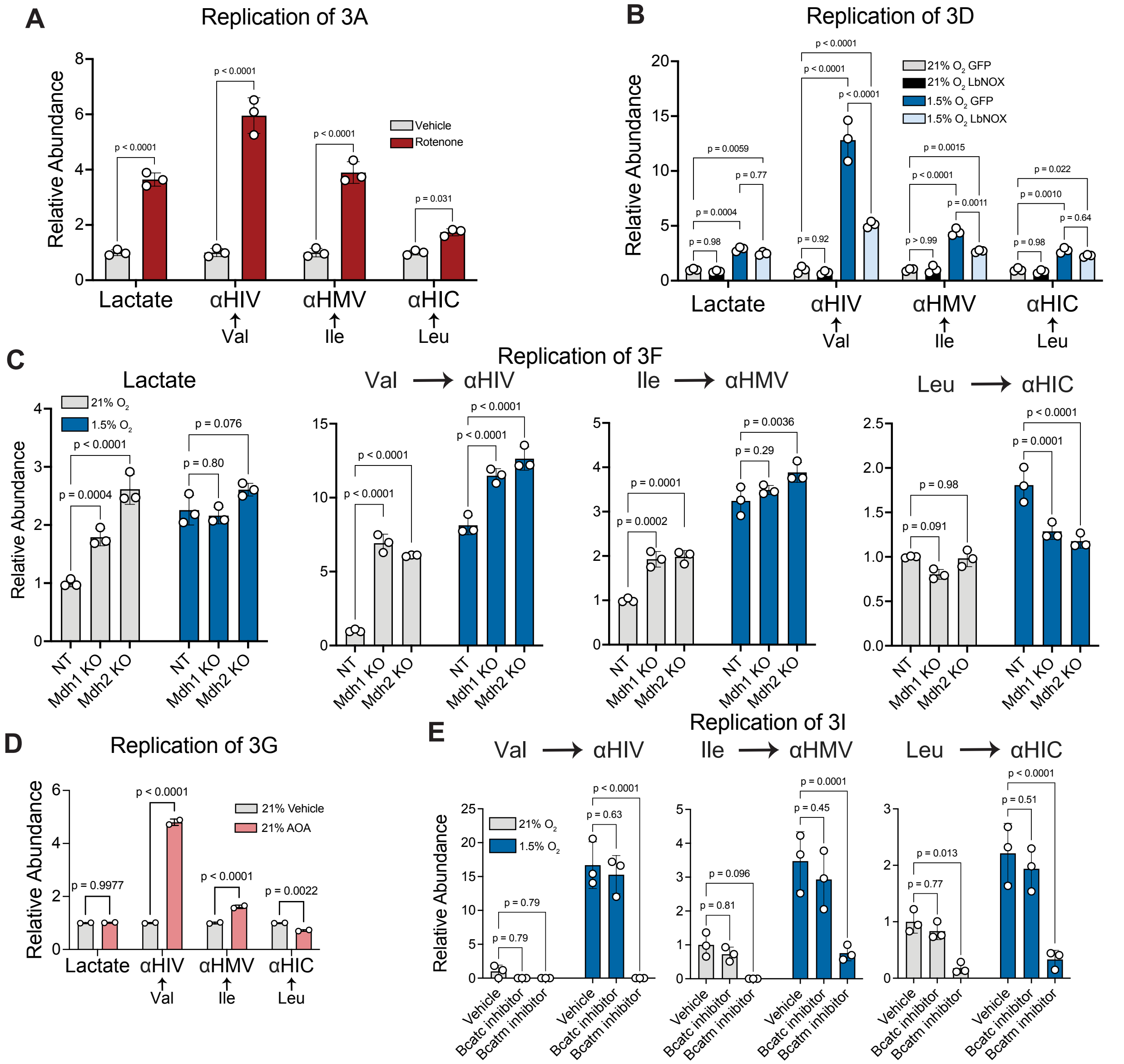


**Figure S5. Compartmental redox signals control BCAA fermentation**

(A) Relative abundance of lactate and $^{13}C$-BCHAs in conditioned media from 3T3-L1 adipocytes treated with 0.1% ethanol or 500 nM rotenone for 3 days. n = 3 wells.

(B) Relative abundance of lactate and $^{13}C$-BCHAs in conditioned media from 3T3-L1 adipocytes after 3-day treatment with 21% $O_2$ or 1.5% $O_2$. n = 3 wells.

(C) Relative abundance of lactate and $^{13}C$-BCHAs in conditioned media from control or *Mdh1*-deficient or *Mdh2*-deficient 3T3-L1 adipocytes after 3-day treatment with 21% $O_2$ or 1.5% $O_2$. n = 3 wells.

(D) Relative abundance of lactate and $^{13}C$-BCHAs in conditioned media from 3T3-L1 adipocytes treated with 400 μM AOA for 3 days. n = 2 wells.

(E) Relative abundance of lactate and $^{13}C$-BCHAs in conditioned media from 3T3-L1 adipocytes after 3-day incubation in 21% $O_2$ or 1.5% $O_2$ and treatment with 10 μM BCATc inhibitor 2 or 50 μM BCAT-IN-2 (BCATm inhibitor). n = 3 wells.

Mean ± SD are shown. Significance was determined by two-way ANOVA followed by Šídák's multiple comparisons test (A, D) or Tukey's multiple comparisons test (B, C, E).

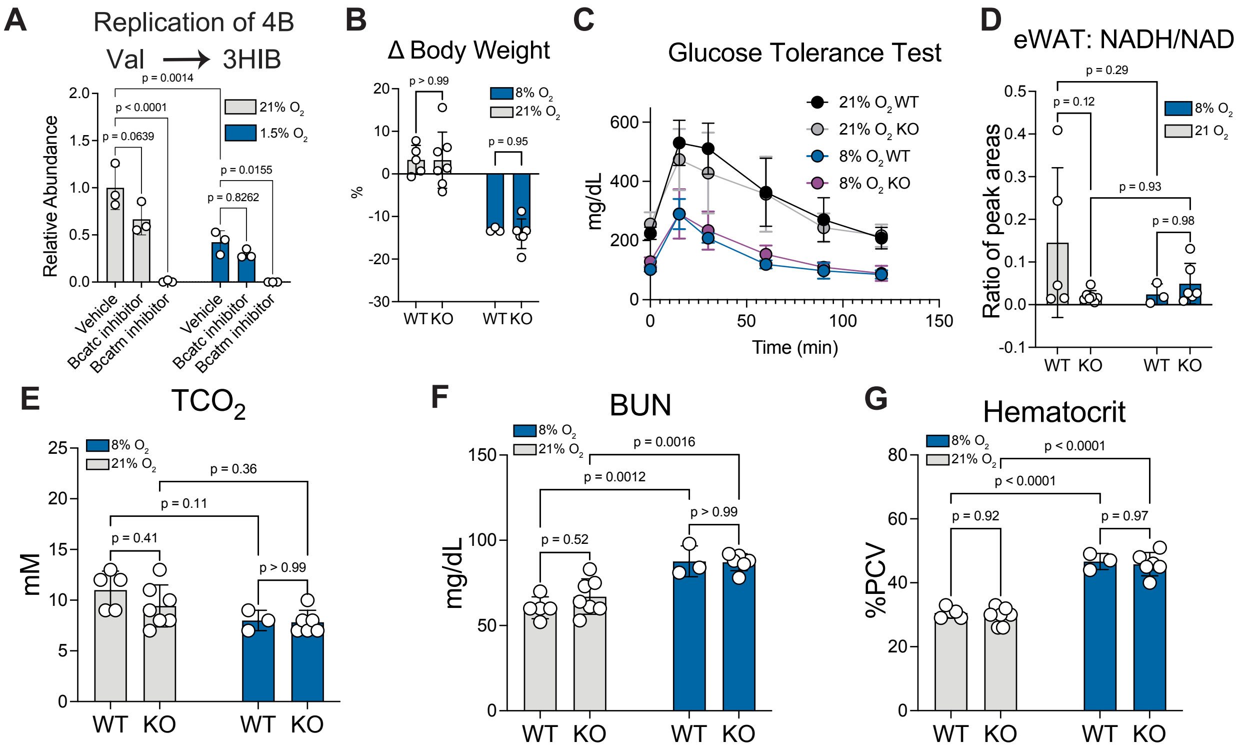


**Figure S6. Phenotyping of *Bckdk*-deficient mice in hypoxia**

(A) Relative abundance of $^{13}$C-3HIB in conditioned media from 3T3-L1 adipocytes after 3-day incubation in 21% $O_2$ or 1.5% $O_2$ and treatment with 10 µM BCATc inhibitor 2 or 50 µM BCAT-IN-2 (BCATm inhibitor). n = 3 wells.

(B) Change in body weight over 12 days at 21% or 8% $O_2$ for wild-type and *Bckdk*-deficient male mice.

(C) Glucose tolerance tests after 12 days at 21% or 8% $O_2$ for wild-type and *Bckdk*-deficient male mice.

(D) Ratio of NADH/NAD$^+$ in eWAT from wild-type and *Bckdk*-deficient male mice housed at 21% $O_2$ or 8% $O_2$ for 3 weeks.

(E-G) Total blood $CO_2$ ($TCO_2$) (E), blood urea nitrogen (BUN) (F), and hematocrit (G) after 12 days at 21% or 8% O2 for wild-type and Bckdk-deficient male mice.

For (B-G), WT/Het n = 5 mice, 21% KO n = 6 mice, 8% WT/Het n = 3 mice, 8% KO n = 6 mice.

Mean ± SEM are shown. Significance was determined by two-way ANOVA followed by Tukey’s multiple comparisons test (A, D-G) or Šídák's multiple comparisons test (B).

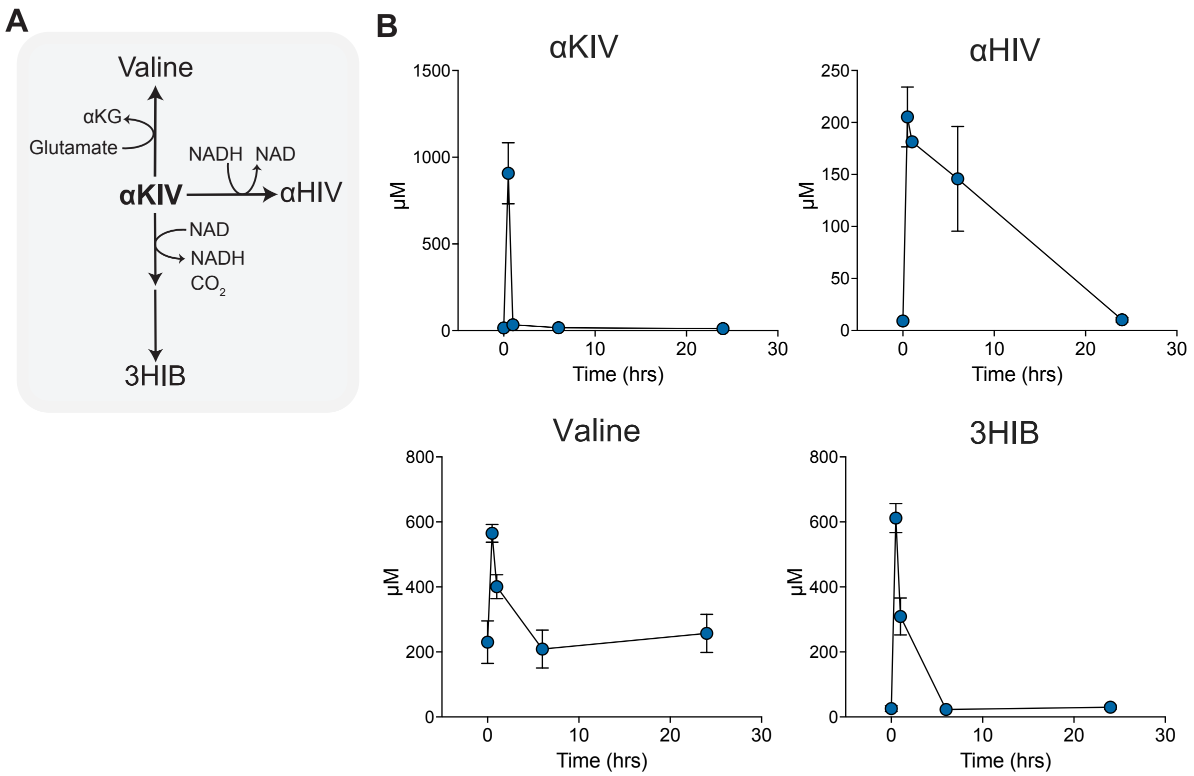


**Figure S7. Dynamics of *in vivo* BCKA supplementation**

(A) Schematic depicting fates of supplemented α-ketoisovalerate (αKIV).

(B) Circulating concentrations of αKIV and its resulting metabolites 30 minutes, 1 hour, 6 hours, and 24 hours after intraperitoneal administration of 1 mg/g of each BCKA. n = 4 mice at time 0, n = 3 mice at 30 minutes, n = 2 mice at 1 hour, n = 3 mice at 6 hours, and n = 5 mice at 24 hours.

Mean ± SD are shown.

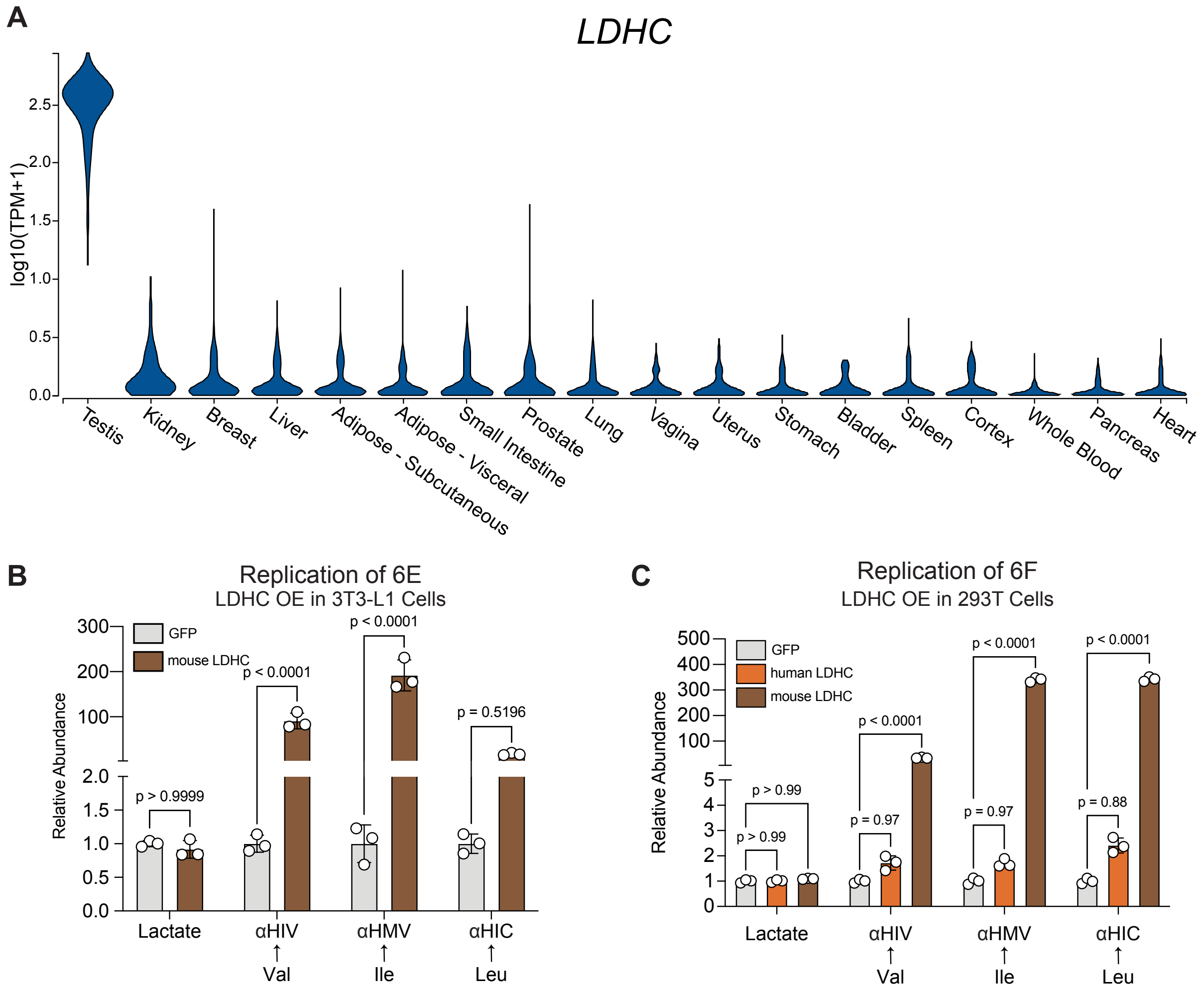


**Figure S8. Lactate dehydrogenase C catalyzes BCAA fermentation**

(A) Bulk gene expression of *LDHC* across human tissue samples from GTEx Analysis Release V10.

(B-C) Relative abundance of lactate and $^{13}$C-BCHAs in conditioned media from undifferentiated 3T3-L1 cells (B) overexpressing GFP or mouse LDHC and from HEK293T cells (C) overexpressing GFP, human LDHC, or mouse LDHC after 3 days of incubation at 21% $O_2$. n = 3 wells.

Mean ± SD are shown. Significance was determined by two-way ANOVA followed by Šídák's multiple comparisons test (B) or Tukey's multiple comparisons test (C).

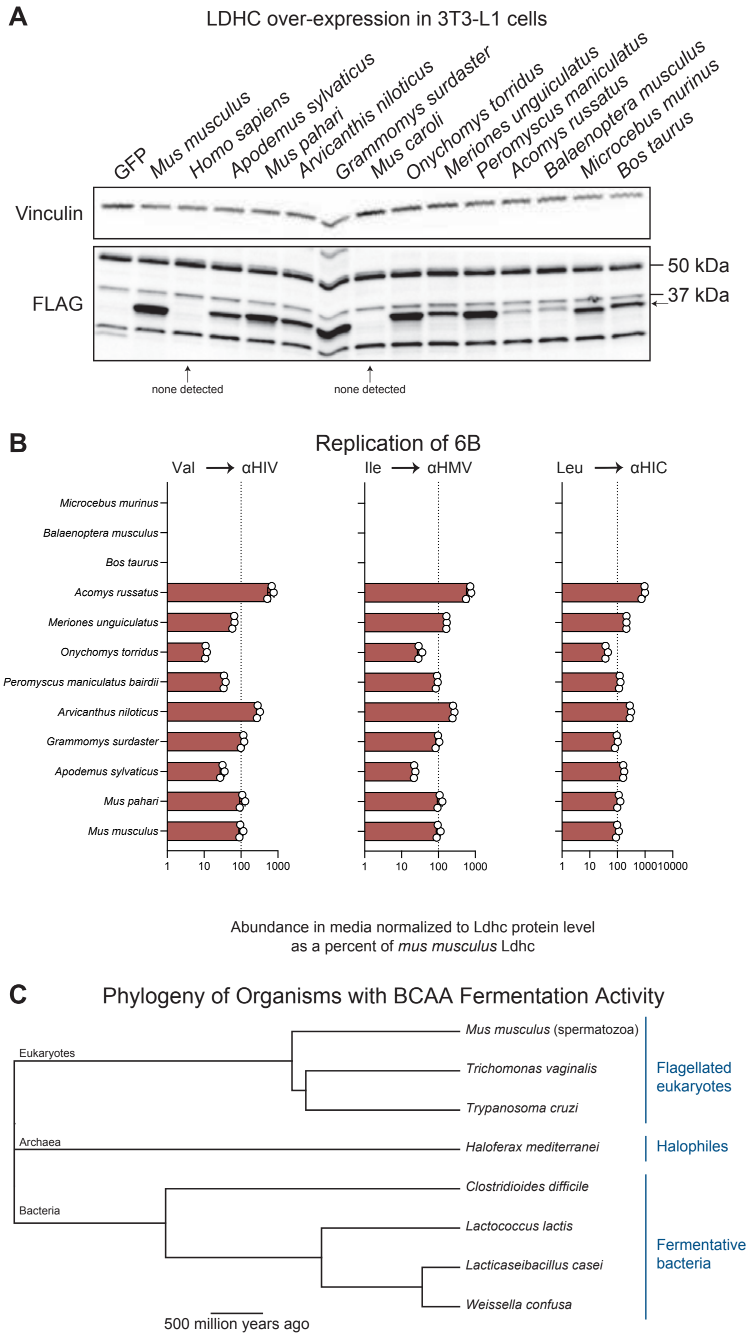


**Figure S9. Expression of LDHC from different mammals**

(A) Immunoblot showing over-expression of FLAG-tagged LDHC proteins from different mammals in 3T3-L1 cells.
(B) Relative abundance of $^{13}$C-BCHAs in the conditioned media from undifferentiated 3T3-L1 cells overexpressing each of the different LDHC sequences. Relative abundances were normalized based on LDHC protein expression, determined by immunoblot, and scaled to the metabolite abundance observed from the cells overexpressing *Mus musculus* LDHC. n = 3 wells for most lines, n = 2 wells for the line expressing *Acomys russatus* LDHC.
(C) Phylogenetic tree of a non-comprehensive list of organisms known to exhibit BCHA dehydrogenase activity.

Mean ± SD are shown.

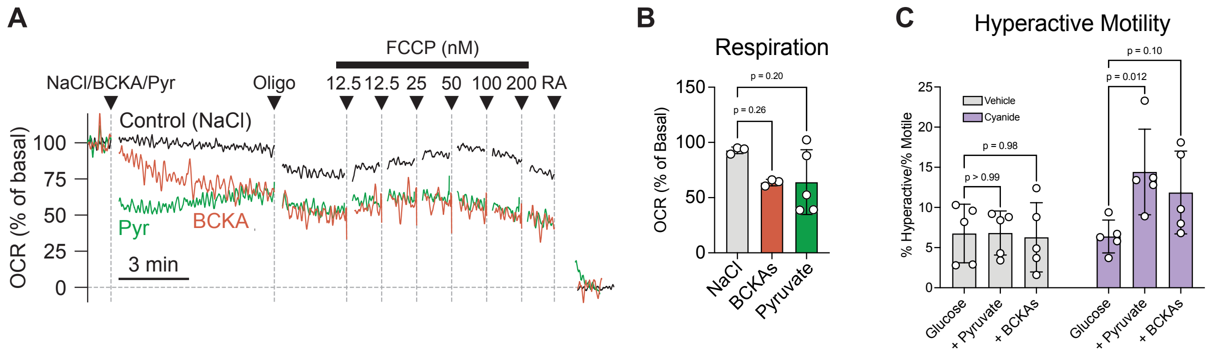


**Figure S10. Respirometry and motility of mouse sperm**

(A) Average oxygen consumption rate (OCR) traces in non-capacitated mouse sperm spiked with 5 mM NaCl, 5 mM BCKA sodium salts, or 5mM sodium pyruvate followed by treatment with oligomycin, FCCP, and rotenone + antimycin.

(B) OCR in non-capacitated mouse sperm spiked with 5 mM NaCl, 5 mM BCKA sodium salts, or 5mM sodium pyruvate as a percentage of basal OCR

(C) Percentage of capacitated mouse sperm motile trajectories that were hyperactive. n = 5 mice. Equal numbers of sperm from each mouse were split into each condition for capacitation.

Mean ± SD are shown. Significance was determined by one-way ANOVA followed by Tukey's multiple comparisons test or two-way ANOVA followed by Tukey's multiple comparisons test (C).
